# Neurons and astrocytes have distinct organelle signatures and responses to stress

**DOI:** 10.1101/2024.10.30.621066

**Authors:** Shannon N. Rhoads, Weizhen Dong, Chih-Hsuan Hsu, Ngudiankama R. Mfulama, Joey V. Ragusa, Michael Ye, Andy Henrie, Maria Clara Zanellati, Graham H. Diering, Todd J. Cohen, Sarah Cohen

## Abstract

Neurons and astrocytes play critical yet divergent roles in brain physiology and neurological conditions. Intracellular organelles are integral to cellular function. However, an in-depth characterization of organelles in live brain cells has not been performed. Here, we used multispectral imaging to simultaneously visualize six organelles – endoplasmic reticulum (ER), lysosomes, mitochondria, peroxisomes, Golgi, and lipid droplets – in live primary rodent neurons and astrocytes. We generated a dataset of 173 Z-stack and 99 time-lapse images accompanied by quantitative analysis of 1418 metrics (the “organelle signature”). Comparative analysis revealed clear cell-type specificity in organelle morphology and interactions. Neurons were characterized by prominent mitochondrial composition and interactions, while astrocytes contained more lysosomes and lipid droplet interactions. Additionally, neurons displayed a more robust organelle response than astrocytes to acute oxidative or ER stress. Our data provide a systems-level characterization of neuron and astrocyte organelles that can be a reference for understanding cell-type-specific physiology and disease.

## Introduction

Multicellular organisms have gained evolutionary advantage through differentiation of specialized cell types. Cells are further compartmentalized into organelles that carry out separate yet coordinated biochemical reactions. Organelles can provide insight into a cell’s physiology and function. This idea is exemplified in specialized cell types such as adipocytes, muscle cells, and neurons, that contain unique organelle landscapes to sustain their dedicated functions. Adipocytes are comprised of a central large lipid droplet for lipid storage and a comparatively smaller cytoplasmic region for other organelles^1^; muscle cells contain an intricate mitochondrial reticulum structure for efficient metabolite diffusion during muscle contraction^2^; and neurons have an abundance of vesicles crowded into presynapses for effective signal transduction^3^. Each of these examples illustrates the importance of a single organelle type to cell function. However, organelles do not operate independently. Organelle interactions at membrane contact sites and areas of close proximity occur to coordinate the activities of individual organelles. Contact sites are involved in signaling pathways and the transfer of ions, lipids, metabolites, and proteins^4^. Moreover, alterations in organelle morphology and interactions are integral to their functionality^5–11^.

Approaches to visualize multivariate phenotypes, such as simultaneous changes in organelle morphology and interactions, can be helpful in inferring organelle and cell state and function. However, techniques to study more than three to four organelles at a time are limited. Electron microscopy (EM) is the most common technique, but it requires harsh fixation protocols. In addition, transmission EM has historically been limited to single thin slices. With the advent of focused ion beam-scanning EM, the field has gained the ability to study volumetric samples, making the analysis of complex cell types such as neurons possible^12^. However, the imaging and analysis procedures are time-consuming and result in low throughput. Recently, our group and others have utilized multispectral confocal microscopy for higher throughput, multi-organelle analysis^13–17^. This approach enables live-cell imaging and results in more accurate population estimations that take cell-to-cell variability into account.

Spectral imaging can provide insight into organelle composition and inter-organelle communication networks. However, a global description of organelles in live primary brain cells, including neurons and astrocytes, has yet to be achieved. Neurons and astrocytes are morphologically and functionally distinct brain cell types. Neurons are specialized to facilitate long-range electrochemical communication and endure over an organism’s lifetime. Astrocytes are a heterogeneous class of neuroglia with many known functions; they regulate synapses, maintain the blood-brain barrier, guide neuronal migration, and regulate immune responses to disease and injury^18,19^. Proteomic characterizations of neurons and astrocytes further reflect their functional specificity, revealing prominent ion binding and receptor and synaptic signaling pathways in neurons, and lipid metabolism and cell-cell interactions in astrocytes^20^. Moreover, in neurodegenerative diseases, including amyotrophic lateral sclerosis (ALS), Alzheimer’s disease (AD), and Parkinson’s disease (PD), pathogenesis is driven by unique functional deficits across multiple cell types, including neurons and astrocytes. These deficits include the dysregulation of diverse cellular processes, from protein quality control systems to mitochondrial dysfunction^21–23^.

Characterizing the landscape of the major membrane-bound organelles involved in these processes holds promise in furthering our understanding of cell-type-specific functions in health and disease. Examination of the endoplasmic reticulum (ER), Golgi (GL), and lysosomes (LS) can provide insight into the secretory pathway^24–27^. The ER is a primary site of lipid synthesis and protein folding; proteins are then sorted at the GL and degraded in LS. Mitochondria (MT), peroxisomes (PO), and lipid droplets (LD) are metabolic organelles that play roles in lipid metabolism^28–30^.

Additionally, MT generate ATP through oxidative phosphorylation^31^, while PO both generate and serve as a sink for reactive oxygen species (ROS)^32^. Multiple organelles, including ER, MT, and LS, also play important roles in calcium signaling^33^. Studying these organelles individually and in combination can paint a holistic picture of a cell’s inner workings.

Here, we present a dataset of three-dimensional (3D) single timepoint and two-dimensional (2D) time-lapse confocal microscopy images containing six simultaneously labeled organelles—ER, LS, MT, PO, GL, and LD—in commonly used primary rodent neuron and astrocyte culture models. The images allow quantitative, single-cell analyses of organelle morphology, inter-organellar interactions, subcellular distribution, and cell morphometrics, resulting in 1418 metrics. Below, we showcase a curated set of metrics to summarize the organelle landscapes of neurons and astrocytes and their responses to acute stress. We highlight key distinctions between cell types and conditions. The entire image and analysis datasets are available to the community through BioImage Archive (http://www.ebi.ac.uk/bioimage-archive) under accession number S-BIAD1445 for further exploration, analysis, and hypothesis generation.

## Results

### Organelle signatures describe the morphology, interactions, and distribution of organelles at the single-cell level

We set out to characterize the variation and responses of organelles that underlie cell type differences in neurons and astrocytes. Our approach utilized quantitative multispectral microscopy of organelles in independent cell culture models (Figure 1A, left)^17,34^. Both cell types were obtained from Sprague Dawley rats, cultures were purified to produce monocultures, and neurons were maintained through synaptogenesis (day *in vitro*; DIV15-16)^35^. The cells were simultaneously labeled with genetically encoded and dye-based organelle markers one day prior to imaging. To visualize the six organelles simultaneously, we utilized a confocal microscope adapted with a spectral detector (Figure 1A, center). At the time of imaging, the organelle labeling process had a minor impact on cell viability and no detectable activation of ER stress from protein overexpression (Figure S1A-B). Upon visual inspection, neurons displayed conical pyramidal shapes without neuritic blebbing or beading^36,37^, astrocytes were elongated with several fine processes each^38^, and nuclei in both cell types displayed normal morphologies (Figure 1B-C, Movie S1)^39^. Both single-timepoint Z-stacks and single-Z-plane time-lapse images were acquired to map the 3D structure of organelles and their dynamics, respectively.

**Figure 1.**
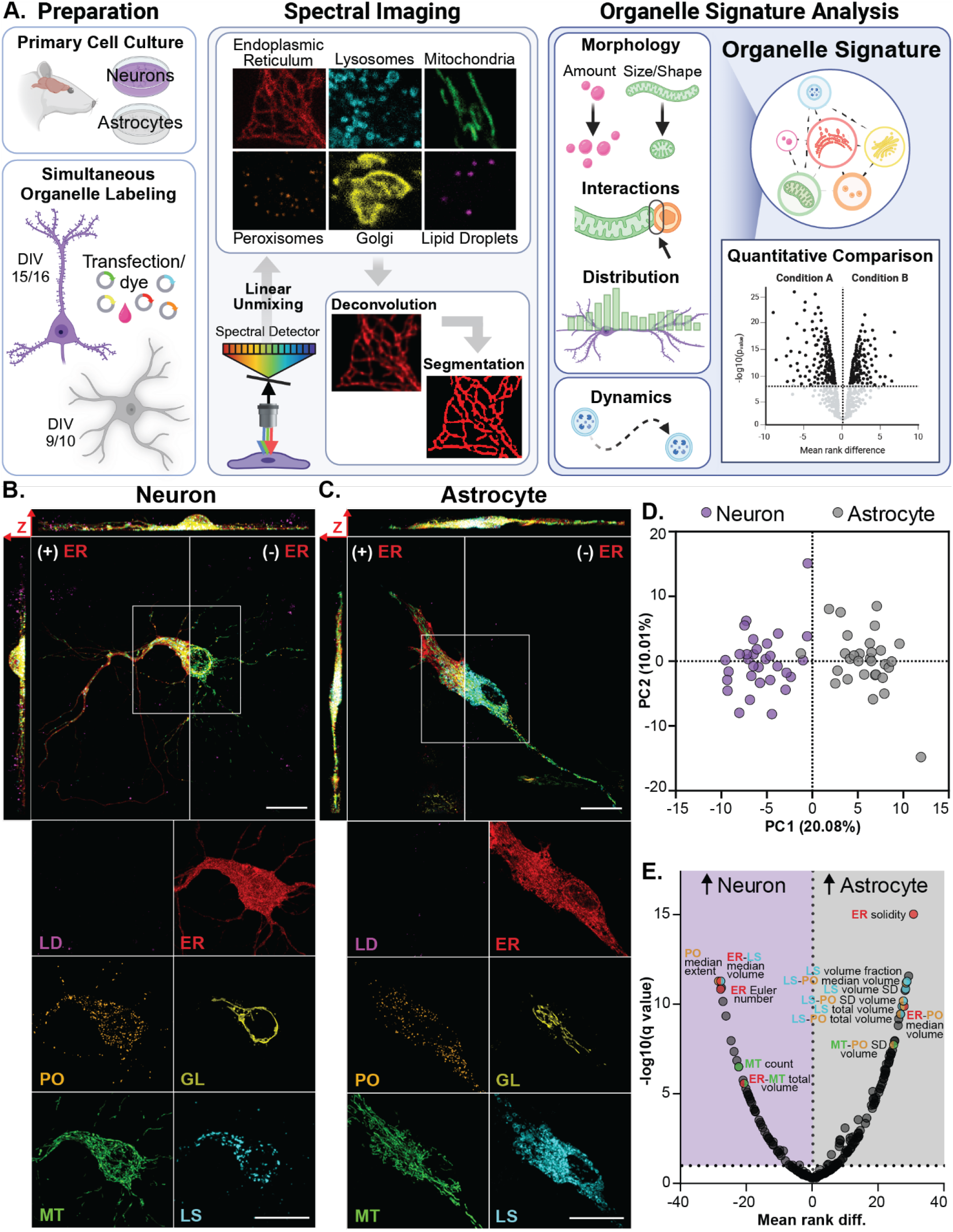
Comparative analysis of neuron and astrocyte organelle signatures. A. Experimental design for multispectral imaging and organelle signature analysis in live primary rat neurons and astrocytes. B-C. Representative multispectral images of a neuron (B) and astrocyte (C) labeled with markers of the endoplasmic reticulum (ER), lysosomes (LS), mitochondria (MT), peroxisomes (PO), Golgi (GL), and lipid droplets (LD). Visualizations are maximum intensity projections across all Z- planes; scale bars represent 20 μm. D. Principal component (PC) scores of neuron and astrocyte organelle signatures; percentages represent percent variance explained. E. Volcano plot of organelle signature metric comparison between neurons and astrocytes. Statistical significance was determined by Mann-Whitney U test with a 10% false discovery rate. Also, see Figure S1 and Table S1.

Spectral images were processed by linear unmixing to produce six-channel images representative of the six organelles (Figure 1A, center). The images were then deconvolved to improve the resolution of fine structures and organelle edges needed for more accurate segmentation. Instance segmentation was performed on each organelle intensity channel from Z-stack images (Figure S1C- D). To enable single-cell analysis, we inferred masks of the cell volume and nucleus for each cell using a combination of intensity channels (Figure S1C-D, right). Segmented organelle objects from each cell were assessed independently for aspects of organelle morphology and subcellular distribution. Then, they were considered in pairwise combinations to quantify organelle interactions. The cell and nuclei masks were also measured for aspects of cellular morphology.

Lastly, organelle dynamics were assessed qualitatively in time-lapse images. In combination, these analyses resulted in 1418 metrics that summarize a particular cell type or cellular condition at a single-cell level. Herein, this summary will be known as the “organelle signature” (Figure 1A, right).

Because organelle signatures are quantitative, comparisons can be made directly across cell types or conditions (Figure 1A, right). The dataset described herein includes two cell types—neurons and astrocytes—and three cellular conditions each: vehicle-treated controls, herein referred to as “controls”, and two acute drug exposures used to induce oxidative and ER stress. These stress conditions perturb the system in a manner associated with neurological disorders where neuron and astrocyte dysfunction are observed^22^. Biological and experimental replicate information is outlined in Figure S1E for all conditions. We have compared organelle signatures across these conditions using a curated set of 234 organelle signature metrics. The curation process was done *a priori* and included consideration for statistical rigor, including limiting metric redundancy and coverage of all possible major phenotypic changes (e.g., size, shape, distribution, interactions, etc.) across each organelle (Figure S1F). This ensured that no one organelle or phenotype was preferentially included based on known cell functions or expected organelle phenotypes within our conditions.

### Neurons and astrocytes display cell-type specific organelle signatures

We first compared the control neurons and astrocytes to establish differences in their baseline organelle signatures. We gained a system-wide perspective on organelle signature differences using a principal component analysis (PCA). Principal component (PC) scores clearly distinguished between the two cell types based on PC1 (Figure 1D). The top metrics contributing to PC1 include size, shape, and interaction differences in LS, PO, MT, and ER (Figure 1E; labels). To elucidate differences metric-by-metric, we conducted a Mann-Whitney U test across all 234 metrics (Figure 1E; Table S1). A false discovery rate (FDR) of 10% resulted in 144 metrics that were significantly different between cell types, while 90 metrics showed no significant difference. These findings are described in more detail in Figures 2-4.

**Figure 2.**
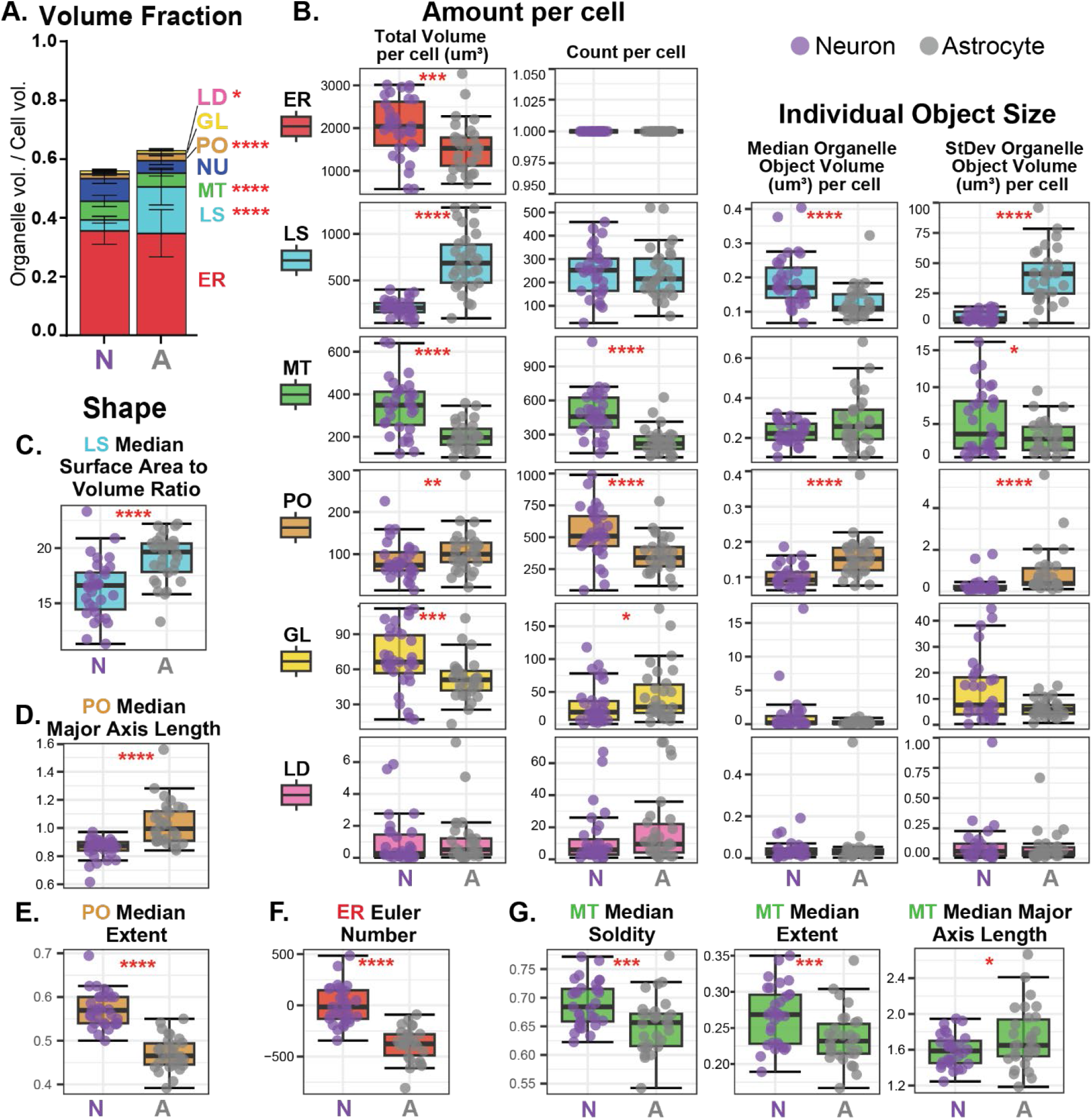
Organelle morphology is cell-type specific. A. Volume fraction of organelles in neurons and astrocytes; bars represent mean ± SD. B-G. Box plots of organelle size and shape metrics per cell; total volume indicates the sum of all the object volumes for the indicated organelle type per cell (B); data points represent single cells. Asterisks (A-G) represent q-values from the combined organelle signature analysis. Also, see Figure S2.

### Organelle abundance relates to neuron and astrocyte functions

We included 65 metrics of organelle morphology (i.e., amount, size, and shape) in our curated analysis (Figure S1F); 34 differed significantly between neurons and astrocytes (Table S1). We chose organelle volume fraction (Figure 2A), total volume, and count (Figure 2B) to assess the amount of each organelle per cell. The volume fraction is the fraction of the cell volume occupied by each organelle type. The proportion of cell volume allotted to each organelle could shed light on the relative importance of that organelle’s function to cellular homeostasis. In contrast, the total volume and count do not consider the cell size but could indicate the total capacity of that cell to carry out organelle-specific functions. All six organelles differed in one or more of these amount metrics between neurons and astrocytes (Figure 2A-B). Generally, neurons contained a higher total volume of ER, MT, and GL, while astrocytes were higher in LS and PO (Figure 2B). The total volume of LD did not differ between cell types, but the volume fraction was higher in astrocytes (Figure 2A). These trends were corroborated by previously published RNAseq data of transcripts encoding canonical organelle proteins from human and rodent brains (Figure S2A)^40,41^.

The most striking difference in organelle amount was a higher LS volume fraction and total volume in astrocytes than in neurons (Figure 2A-B). Supporting our findings, the canonical LS protein LAMP1 transcript was reported to be more highly expressed in astrocytes (Figure S2A)^40,41^. This data implies that astrocytes have a higher lysosomal capacity and require more lysosomal activity for homeostasis than neurons. Conversely, higher MT volume fraction, total volume, and count in neurons indicate that mitochondrial functions, such as ATP production through oxidative phosphorylation (OXPHOS), may be needed at a higher capacity to achieve proper neuronal homeostasis. Consistent with this observation, the transcripts for the inner mitochondrial member protein and respiratory chain component Cox5a and the outer mitochondrial membrane protein Tomm20 were reportedly higher in neurons (Figure S2A)^40,41^.

### Organelle morphology is unique between cell types

Our curated analysis also included measurements of organelle object size—individual object volume and standard deviation (SD) of object volumes per cell—and shape—surface area to volume ratio, equivalent diameter, extent, Euler number, solidity, and major axis length. The median value per cell for each metric, except the SD of volumes per cell, was included in our curated analysis. We identified nuanced differences in the size and shape of each organelle between cell types except LD (Table S1), which were spherical and relatively low in abundance (Figure 1B-C, S1C-D, 2B).

In astrocytes, the LS population had a lower median object volume but a higher SD of volumes per cell (Figure 2B). Because segmented organelle objects in our analysis can represent closely clustered groups of organelles as well as individual organelles, these metrics indicated that astrocytes either contained a wide range of relatively spherical LS objects or a variety of different-sized LS clusters. We examined differences in LS object shape to better understand which phenotype was occurring. Most notably, the median LS surface area to volume ratio was higher in astrocytes (Figure 2C). Since the surface area to volume ratio of a sphere reduces as volume increases, it is more likely that astrocytes contain clusters of LS. Example images confirmed these findings (Figure 1A-B). We also note that individual LS vesicles in intensity images appeared larger in astrocytes than in neurons.

Distinct from LS, the PO population in neurons had a higher count and a lower median and SD of volumes per cell (Figure 2B). This represents a smaller but more numerous PO phenotype in neurons. We also observed that most PO shape measurements differed between cell types (Table S1). In particular, the PO major axis length, the longest distance across an ellipsoid of the same second central moment as the object (Figure S2B), was higher in astrocytes (Figure 2D), suggesting that astrocytes have longer PO objects than neurons. We then looked to the extent metric, or the amount of the bounding box filled by the object, to better understand if the objects were more spherical or elongated (Figure S2C). The median extent value was higher in neurons (Figure 2E), indicating the objects were more spherical. Representative images in Figure 1 exemplify these phenotypes, showing that neurons contained more numerous but smaller PO that were not clustered into groups, while astrocytes had larger PO objects that would occasionally form small, elongated clusters (Figure 1B-C and S1C-D). Interestingly, PEX11β, a peroxisomal protein with higher transcript levels in neurons (Figure S2A)^40,41^, was reported to increase peroxisomal proliferation rates^42,43^. This could explain the presence of more numerous smaller peroxisomes in neurons. A loss of PEX11β has also been implicated in peroxisomal clustering^44^. Lower PEX11β transcript levels could indicate the role of this protein in astrocyte PO clustering.

### Some organelle morphology phenotypes correlate with cell morphology

Many other size and shape differences were evident in our curated dataset (Table S1), including differences in more morphologically complex organelles such as the ER and MT. The Euler number, a measure of object topology that considers the number of surface holes and tunnels through an object (Figure S2D), is higher in neurons (Figure 2F). This finding indicates the ER in neurons contains fewer tunnels than in astrocytes. Representative maximum intensity projections confirm this finding (Figure 1B-C). Moreover, neuronal MT had a higher median solidity and extent and lower major axis length than astrocytic MT (Figure 2G and S2E). Together with the higher SD of MT volumes in neurons (Figure 2B), these metrics indicated a wider range of MT sizes in neurons, possibly including more fragmented MT.

Because these organelles, especially the ER, can create a connected network throughout the cell, we wanted to assess the correlation between cell morphology and organelle morphology. We performed a correlation analysis of organelle signature metrics to cell morphometrics for both cell types (Figure S2F, Table S2). Some organelle metrics, including the count and total volume of LS, MT, PO, and ER, positively correlated with metrics of cell size (e.g., volume, equivalent diameter) in neurons and astrocytes. This finding indicates that some organelle amounts are scaled to cell size. However, most organelle shape metrics had a much weaker correlation with cell size or shape, and some even had negative correlations. Some exceptions included the ER equivalent diameter and extent, which significantly correlated with several cell metrics, including cell volume, equivalent diameter, and extent. This finding was not surprising since the ER network traverses most of the cell volume but indicates that ER equivalent diameter and extent differences between cell types are likely due to differences in neuron and astrocyte cell shape, not organelle shape.

### Distinct organelle interactomes and interaction dynamics reflect functional and metabolic differences between neurons and astrocytes

Next, we examined organelle interactions in neurons and astrocytes by quantifying the amount and size of regions of high proximity between pairwise combinations of organelles, herein known as organelle interaction sites. These interaction sites were defined as locations where two segmented organelle objects appeared less than one voxel’s distance apart in any direction (Figure S3A). At our imaging resolution, these sites included membrane contact sites (10-80 nm) and areas of close proximity (<410 nm). The total volume, count, median object volume, and SD of volumes per cell for all 15 pairwise interaction sites were included in our curated analysis (Table S1), resulting in 60 interaction metrics (Figure S1F). Together, these metrics define the organelle interactome.

Based on interaction site counts, the relative abundance of organelle interactions was similar between cell types: ER interactions were the most abundant, followed by MT, PO, LS, GL, then LD (Figure 3A). As expected, the highest-volume organelle, ER, and the lowest-volume organelles, GL and LD, had the highest and lowest total interaction volumes, respectively (Figure 3A). However, we also identified 42 organelle interaction metrics that significantly differed between neurons and astrocytes, likely reflecting their differing cellular functions and metabolisms (Table S1).

**Figure 3.**
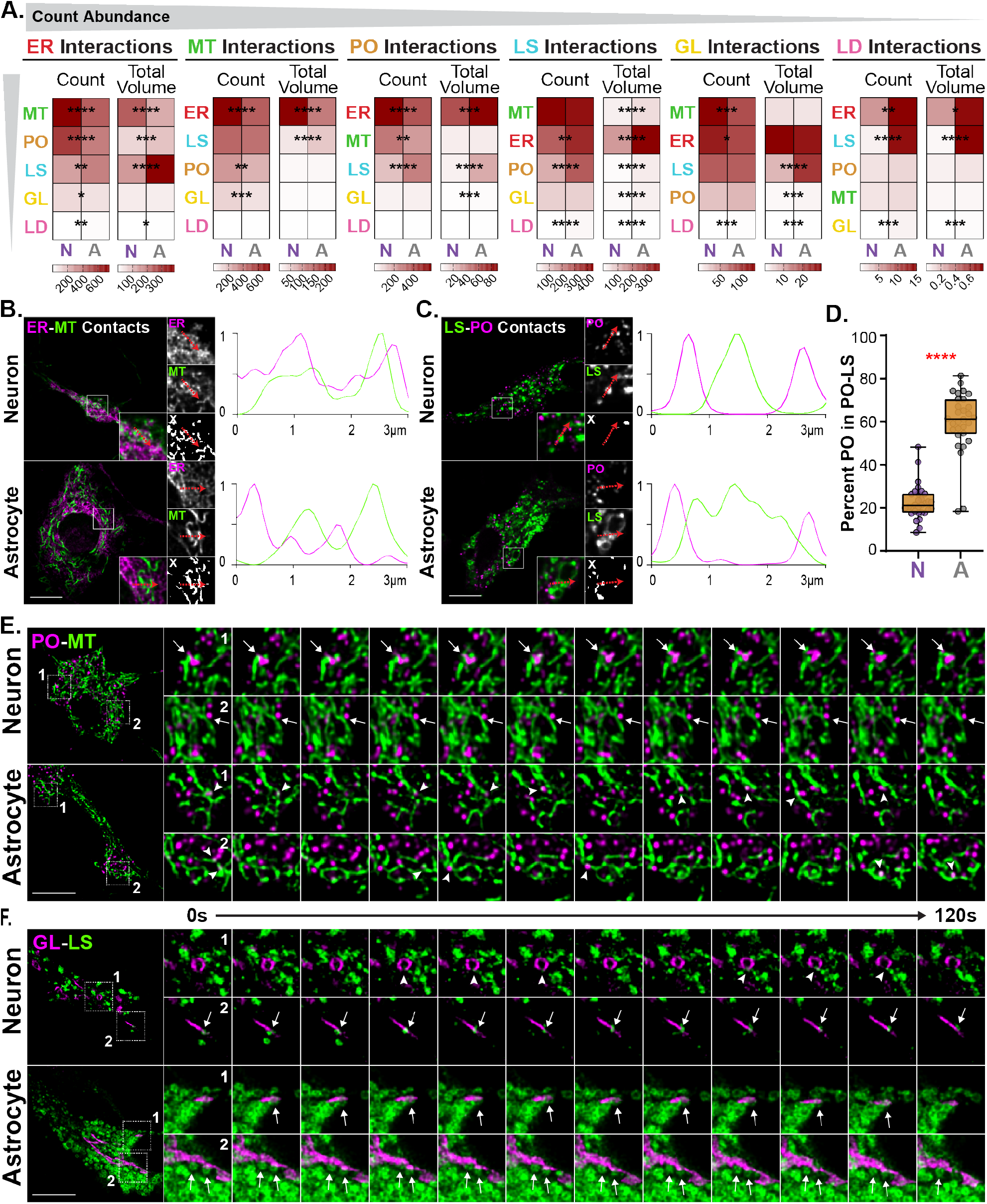
Distinct organelle interactomes and interaction dynamics in neurons and astrocytes. A. Heatmaps displaying the average organelle interaction site count per cell and total volume (μm^3^) per cell of all pairwise organelle combinations listed in order of count abundance; intensity scales under each heatmap differ based on the relative amount of interaction count or total volume per organelle type; asterisks denote q-values from combined organelle signature analysis. B-C. Representative intensity images and interaction sites (X) of ER-MT (B) and LS-PO (C) interactions in neurons and astrocytes; scale bars are 10 μm. Line scan graphs display the normalized intensity measured across the red arrows shown. D. Box plots of the percentage of PO interacting with LS; data points represent single cells; asterisks represent p-value determined by independent t-test. E- F. Representative intensity frames from timelapse images; frames are 10 seconds apart; scale bar is 10 μm. Arrows indicate constant interactions over the 5-minute imaging duration; arrowheads indicate transient interactions. Also, see Figure S3.

Neurons contained more ER-MT interactions with significantly higher metrics of amount and size compared to astrocytes (Figure 3A; Table S1). Within the neuron ER-MT interaction sites, the organelles appeared more closely intertwined, indicated by the regions of overlap that appear white in the merged intensity images or as shared area under the intensity curves in the example line scan (Figure 3B). PO-MT, MT-GL, and ER-GL interactions were also more common in neurons. These three interaction types were higher in count in neurons but differed in which size metrics were distinct between cell types (Figure 3A; Table S1). PO-MT sites were smaller but more numerous in neurons; MT-GL sites were more numerous in neurons but had a wider range of sizes in astrocytes; and ER-GL sites were the same size and volume in both cells but occurred slightly more often in neurons. These differences could indicate nuances in organelle co-regulation. However, the implication of different organelle interaction site sizes and shapes is not yet well defined.

In contrast, astrocytes had more prominent lysosomal interactions. For example, LS-PO and LS-LD interactions were higher in all four measurements of interaction amount and size (Figure 3A; Table S1). Within the astrocyte LS-PO interaction sites, PO appeared closer to LS, as exemplified by the distance between peaks in the line scan analysis (Figure 3C). It was also evident in example images that a larger fraction of the astrocyte PO population was involved in LS-PO interactions (Figure 3C). We revisited the larger dataset of 1418 metrics to find that the percentage of PO objects involved in LS-PO interactions was significantly higher in astrocytes (Figure 3D). We also found higher levels of LS-LD interactions in astrocytes, but in our conditions, neurons had an average LD count of 12.29 compared to astrocytes with 18.53 (Table S1), making differences in LD interactions statistically more likely in astrocytes and, therefore, challenging to interpret.

Leveraging time-lapse images, we further explored two interaction types: PO-MT and LS-GL. Our time-lapse images contained single Z-plane snapshots every five seconds over a five-minute duration. PO in neurons regularly maintained interactions with MT over the duration of the imaging period (Figure 3E, arrows; Movie S2), while PO only transiently interacted with MT in astrocytes (Figure 3E, arrowheads; Movie S2). This suggests that neurons contain more numerous PO-MT interaction sites due to the higher retention rate of the interactions over time. LS-GL interactions, which were higher in total volume, median volume, and SD of volumes in astrocytes (Figure 3A; Table S1), were sustained throughout the imaging period in both neurons and astrocytes (Figure 3F, arrows; Movie S3). However, in neurons, LS-GL interactions occurred less frequently and seemed to happen preferentially at the endpoints of the GL objects. Transcripts for the tether proteins Rab34 and FLCN, which engage to form LS-GL membrane contact sites through the effector protein RILP^45^, were reportedly higher in astrocytes, corroborating our findings (Figure S3B)^40,41^. In astrocytes, we also qualitatively observed reduced LS motility in regions surrounding the GL where LS-GL interactions occurred regularly, as previously reported^45,46^. LS subpopulation differences in dynamics were not evident in neurons where LS-GL interactions were less prevalent.

### Organelles have distinctive distributions in XY and Z

The subcellular distribution of organelles has been shown to have important impacts on organelle function^47^. Here, we profiled and compared the subcellular distribution of organelles and organelle interactions in neurons and astrocytes (Figure 4). Our curated analysis included 109 metrics from subcellular distribution measurements in XY and Z, independently, for all six organelles, 15 pairwise interaction sites, and the nucleus (Figure S1F). The XY distribution describes the spread of organelles from the nucleus to the cell membrane, while the Z distribution quantifies the spread of organelles from the bottom to the top of the cell (Figure S4A). The cytoplasm from the center of the nucleus to the plasma membrane was divided into five concentric XY regions where the nucleus is included in region one, and the neurites are included in region five (Figure S4A, bottom). Because the size of neurons and astrocytes in the Z dimension was variable cell-to-cell and neurons and astrocytes were differing heights (Figure S4C), Z slices were grouped into ten approximately equal regions per cell, with the bottom of the cell included as region one and the top of the cell as region ten (Figure S4A). The volume of organelles, interaction sites, and the nucleus mask were normalized to the cell and total object volume and measured per XY and Z region (Figure S4B-D).

**Figure 4.**
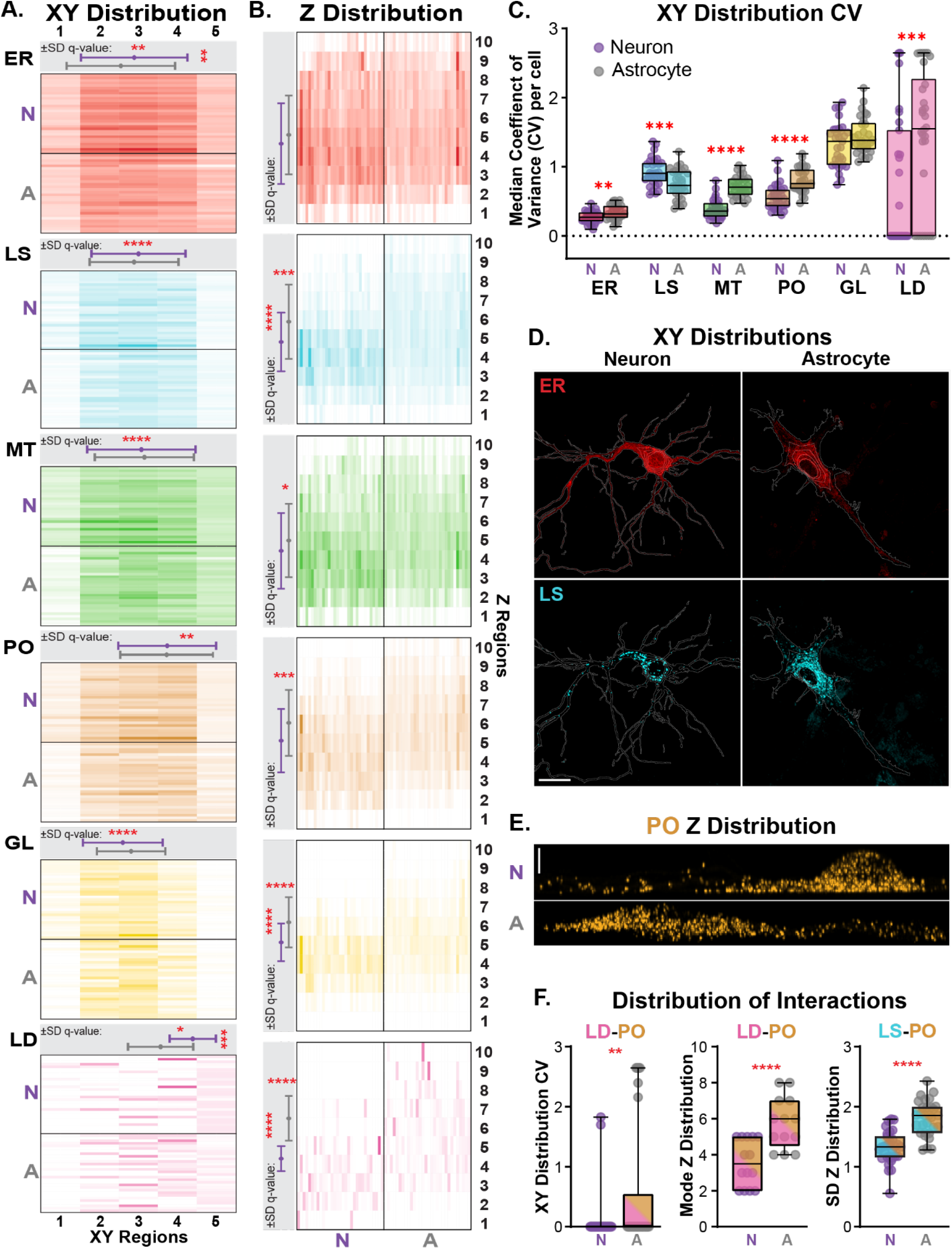
Subcellular distributions of organelles and organelle interaction sites. A-B. Heatmaps of the normalized organelle volume per XY (A) and Z (B) regions; XY region 1 (nucleus), XY region 5 (periphery), Z region 1 (bottom), Z region 10 (top). Grey boxes summarize the mode (data point; asterisk perpendicular to the bars) and SD (error bars; asterisk parallel to the bars) distribution metrics per organelle. Rows (XY) and columns (Z) represent individual cell values per region. C. Box plots of the XY distribution CV metric; individual data points are median CV values per cell. D. Maximum projections of representative ER and LS intensity images overlayed on the XY distribution regions (outlines); scale bar is 20 μm. E. Maximum projections of presentative PO intensity images; image orientation displays the XZ-axes; scale bar is 5 μm. F. Box plots of distribution metrics for organelle interactions; data points represent per cell values. Asterisks (A, C, F) represent q-values from the combined organelle signature analysis. Also, see Figure S4.

Two histogram summary statistics, mode and standard deviation (SD), were included in the curated analysis to succinctly summarize these values for each cell in XY and Z separately (Figure 4A-B, grey boxes; Table S1). The coefficient of variance (CV) within each XY region was also measured to assess the variation of organelle and interaction site volumes per concentric region; the median CV value per cell was included in the curated analysis (Figure 4C).

The perinuclear to peripheral (XY) distribution of organelles was similar between neurons and astrocytes (Figure 4A). The mode, or the XY region with the highest organelle volume, and the SD, or spread of the organelles across the five XY regions, were stereotyped per organelle (Figure 4A, grey box). The GL was perinuclear, the ER, MT, and LS were centrally localized between the nucleus and plasma membrane, and PO and LD were more peripherally localized. Notably, subtle yet significant differences in ER and LD distributions were seen between cell types. ER and LD were more peripherally localized, with a lesser SD across the regions in neurons. Additionally, the ER and LD CV values were lower in neurons (Figure 4C), indicating more consistency in volumes across each of the five concentric XY regions. Though less exaggerated in some cells, example images of the ER clearly portray a less densely packed peripheral region in astrocytes than neurons (Figure 4D, top). Fitting with our findings, the ER is known to play many important roles in the axons and dendrites of neurons and has been shown to maintain a complex structure within these regions^48^. The median CV values for PO and MT were also higher in astrocytes, while the LS median CV value was higher in neurons. Utilizing the LS as another example, we saw that there was a clear difference in organelle volume per region that was distinct from the ER CV phenotype. Neurons contained a lower density of organelles throughout the somatodendritic region than astrocytes, which contained a very dense perinuclear region and a more sparsely populated peripheral region (Figure 4D, bottom). LS perinuclear clustering has been linked to differences in nutrient states^49^, suggesting neurons and astrocytes may contain different metabolic programming in our culture system.

The spread of organelles from the bottom to the top of the cell was variable between neurons and astrocytes, with only the ER showing no statistical differences (Figure 4B). All other organelles had a lower mode in neurons (Figure 4B, grey box), indicating the organelles were localized closer to the bottom of the cell. The PO exemplify this phenotype well (Figure 4E). Additionally, the SD of GL, LS, and LD was greater in astrocytes, suggesting these organelles were inhabiting a wider Z range.

These findings contrasted with the Z distribution of the nucleus (Figure S4D), which displayed only a modestly higher Z mode in neurons. Together, our data shows that cultured neurons maintain their organelles at a lower Z level than astrocytes without a drastic difference in nucleus localization.

### Subtle differences in interaction site distributions exist between neurons and astrocytes

The functional implications of distribution differences in organelle interactions have not yet been explored in depth. However, our analysis included these metrics to begin classifying differences across cell types (Figure 4F; Table S1). Our analysis determined that 45 interaction site distribution metrics were significantly different between cell types. Since organelle distribution can impact interaction distributions, we performed a correlation analysis to determine which organelle interactions were influenced by their constituent organelles (Table S2; Figure S4E). Only three of the significant interaction metrics—LD-PO CV, LD-PO Z mode, and LS-PO Z SD—were not concomitantly correlated with their constituent organelle metrics (Figure 4F). These findings indicate that LD-PO interaction volumes were more variable across XY regions and localized towards the top of the cell in astrocytes. LS-PO interactions were distributed across a larger Z region in astrocytes. The physiological significance of these findings is currently unclear.

### Organelle signatures respond to acute stress in a cell type- and stress-specific manner

An important aspect of cellular physiology is the ability of a cell to respond to environmental cues and maintain homeostasis. This involves the coordination and regulation of organelles. We explored how neurons and astrocytes modify their organelle signatures in response to acute drug exposure. We selected two stress conditions commonly implicated in neurodegenerative diseases: oxidative and ER stress^50^. We examined their impact on organelles using the same 234 curated organelle signature metrics outlined in Figures 1-4.

Oxidative stress was induced with a one-hour 50 μM sodium arsenite exposure, sufficient to induce stress granule formation (Figure S5A), an established cytoplasmic reorganization following arsenite treatment^51^. ER stress was induced with a one-hour 25 nM thapsigargin exposure, which initiated the unfolded protein response (UPR)^52^, evidenced by the presence of phosphorylated eIF2α (Figure S5B). Both conditions were sublethal at the one-hour imaging time point as assessed by measures of lactate dehydrogenase levels and had no lasting impact on cell viability 24 hours after the one-hour exposure by measures of cellular metabolic activity (Figure S5C-D).

Neuron and astrocyte organelle signature changes are displayed in Figures 5A-B, respectively, and summary statistics for each condition are included in Table S3. Heatmaps represent the log_2_(fold change) from the control baselines previously described (Figures 1-4; Table S1). Organelles in both cell types responded to arsenite exposure, but only neurons showed significant organelle differences after thapsigargin treatment. Additionally, the arsenite-induced changes were more numerous than thapsigargin-induced changes. Notably, most of these occurred in organelle interaction metrics, not organelle morphology or distribution metrics. This is likely because organelle interaction changes can be modulated rapidly by specific post-translational modifications^53^, while changes in organelle amount require transcriptional programs to induce organelle biogenesis or turnover^54–57^.

**Figure 5.**
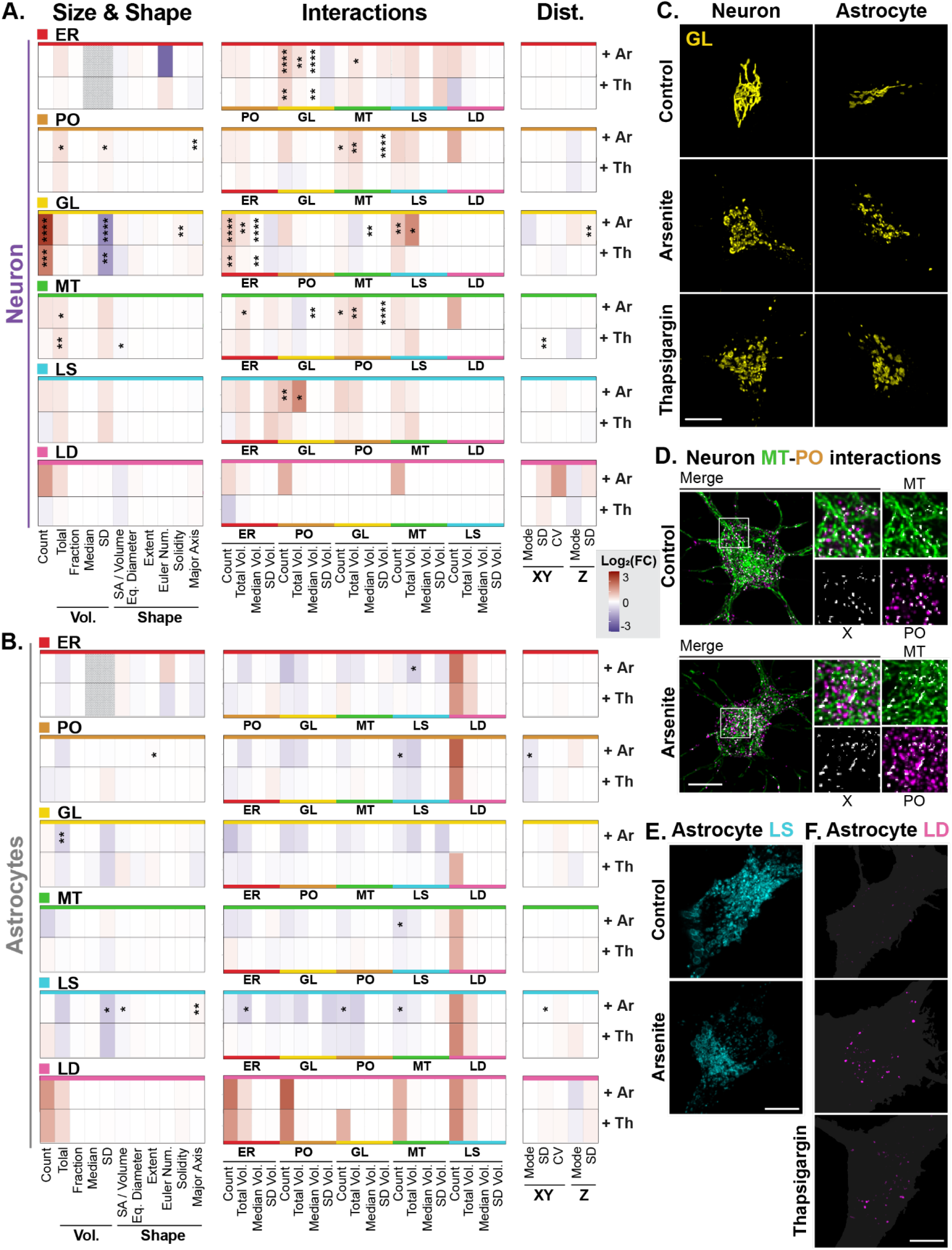
Neuron and astrocyte organelle signatures elucidate differential responses to acute oxidative and ER stress. A-B. Heatmaps of the organelle signature metrics log_2_(fold change) from baseline upon 50 μM sodium arsenite (Ar) or 25 nM thapsigargin (Th) exposure for 1 hour; asterisks denote q-values from combined organelle signature analysis. C-F. Representative maximum intensity projections of GL from neurons and astrocytes in control or drug-treated conditions (C), MT and PO from neurons overlayed with MT-PO interaction sites (X) with and without sodium arsenite exposure (D), and LS and LD from astrocytes with and without drug exposures (E-F); scale bars are 10 μm. F. Cell mask represented as a grey background. Also, see Figure S5 and Table S3.

Oxidative and ER stress share similarities in their mechanisms^58^. Likewise, we found similarities between arsenite- and thapsigargin-induced oxidative and ER stress in our organelle signature dataset. Specifically, GL morphology metrics trended similarly in response to both drug exposures across cell types (Figure 5C). This shared phenotype was most strongly seen in neurons, where our data showed an increase in GL count and a reduction in SD of volumes (Figure 5A), indicating a shift to more uniformly sized Golgi fragments. Our findings corroborate previous reports that GL fragmentation is a common stress response following oxidative and ER stress conditions^59–61^.

Despite this commonality, there are also many established mechanisms unique to arsenite or thapsigargin, especially at early time points^62,63^. Our dataset similarly revealed organelle distinctions between arsenite and thapsigargin responses. For example, increased MT-PO interactions were unique to arsenite-exposed neurons (Figure 5A, D). We found an increase in MT- PO count, total volume, and SD of volumes (Figure 5A). This phenotype was coordinated with a change in PO object morphology—increased total volume, SD of volumes, and major axis length (Figure 5A)—reminiscent of the PO clustering phenotypes in control astrocytes. Coordination of MT and PO at interaction sites and increased PO clustering could reflect compensatory mechanisms to reduce ROS levels following arsenite damage to MT that is specific to neurons^64,65^.

Interestingly, many of the neuron phenotypes, including increased MT-PO interactions, were not evident in astrocytes despite equal treatment concentrations and exposure times. A qualitative comparison between neuron and astrocyte stress signatures revealed opposing trends in most metrics (Figure 5A-B). Astrocyte-specific stress signatures were linked to reduced LS SD of volume per cell and interactions, especially in the arsenite condition, and increased LD amount and interactions trended across both drug exposures (Figure 5B, E-F). Neither of these phenotypes was prominent in neurons, suggesting organelle signature remodeling is strongly cell-type-specific in the face of acute oxidative and ER stress.

### Stress-induced organelle modifications were subtle compared to cell type differences

Lastly, we compared all six drug-by-cell-type conditions. We performed a PCA using the 234 curated organelle signature metrics for all data discussed in Figures 1-5 and plotted each cell based on their top two PC scores (Figure 6A). Neurons and astrocytes maintained a clear separation based on PC1, similar to the results from Figure 1D. These cell-type differences were evident irrespective of drug exposure condition. However, the subtle differences between drug exposure conditions within each cell type remained discernable from the same components (Figure S6A-B). Analysis of the PC scores revealed shifts in the drug treatment populations along the PC2 axis. Specifically, neurons treated with arsenite and, to a lesser extent, thapsigargin had higher average PC2 scores. This contrasted with astrocytes exposed to arsenite, which had lower average PC2 scores. Additionally, neurons and astrocytes displayed opposing shifts in the average PC1 variable during thapsigargin exposure (neuron: 5.11 to 6.67; astrocyte: -5.46 to -6.43). Though subtle (Figure 6B), these differences exemplify the importance of cell type background in understanding cellular responses to perturbations.

**Figure 6.**
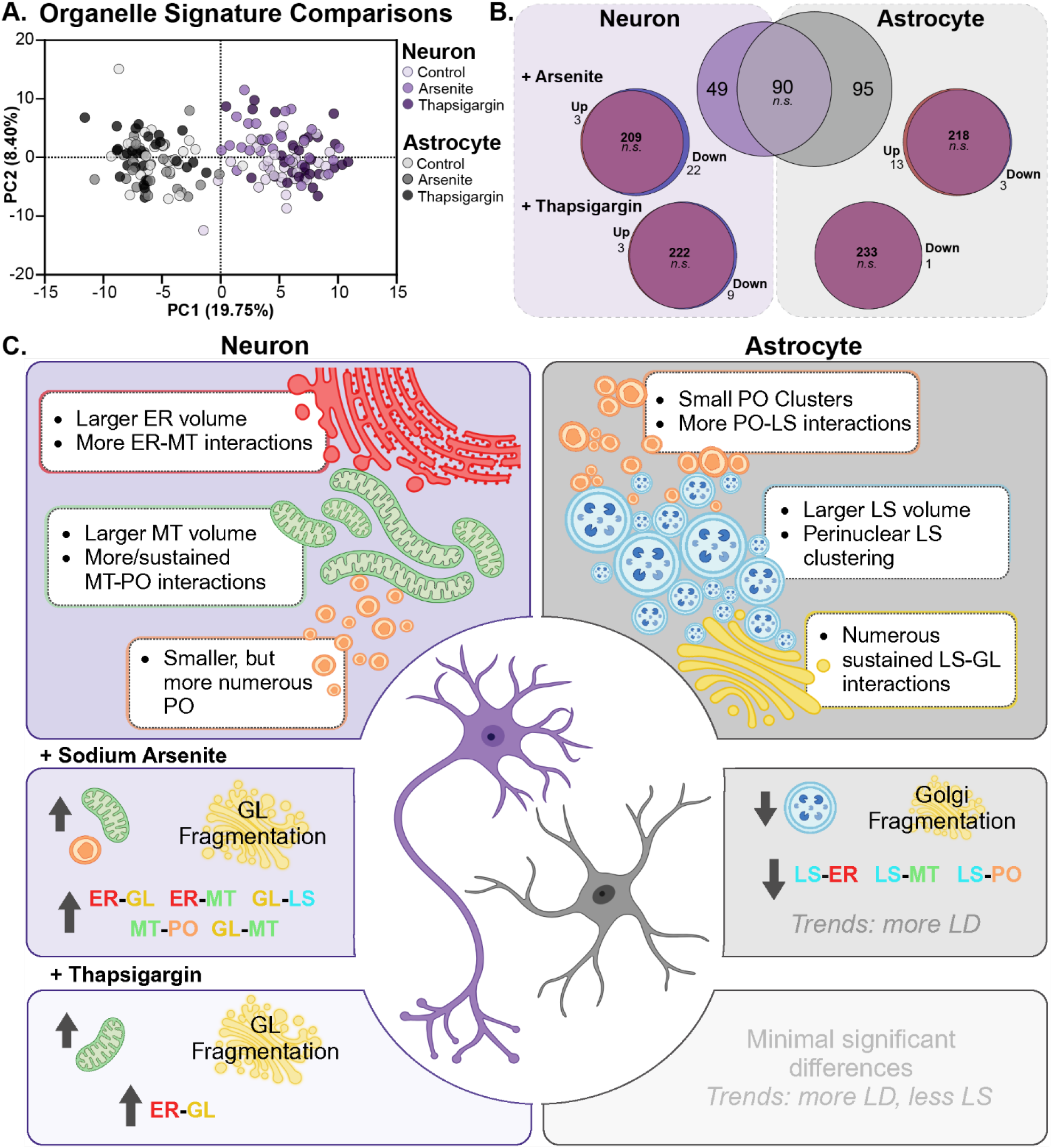
Stress-induced organelle modifications are subtle compared to cell type differences. A. PC scores of all neuron and astrocyte conditions; data points represent individual cells; percentages indicate the percent variance explained per PC. B. Numbers of not significant (*n.s.*) versus significantly different organelle signature metrics between conditions explored in Figures 1-5. C. A summary of neuron and astrocyte organelle signature phenotypes exemplified in the curated analysis at baseline (Figure 1-4) and in response to oxidative and ER stress (Figure 5). Also, see Figure S6.

Hierarchical clustering also revealed unique trends within our dataset (Figure S6C). Again, we could separate neuron and astrocyte populations, irrespective of drug exposure. However, this analysis did not demonstrate a clear clustering of cells based on drug exposure within each cell type. It also revealed no large experiment-to-experiment batch effects in organelle signature metrics, confirming the robustness of our dataset.

## Discussion

Our study presents the first systems-level characterization of organelles in live primary rodent neurons and astrocytes. Our dataset includes 173 Z-stack and 99 time-lapse confocal microscopy images of six major membrane-bound organelles—ER, LS, MT, PO, GL, and LD—from each cell. We quantified 1418 metrics of organelle morphology, interactions, and distribution per cell, enabling us to quantitatively compare across cell types and conditions. A curated list of 234 metrics was sufficient to distinguish between cell types and elucidate nuanced alterations in organelles following acute stress (Figure 6C). Neuron organelle signatures were higher in MT amount and MT interactions with ER and PO. In contrast, astrocyte organelle signatures contained more LS that were clustered in the perinuclear region. LS interactions with GL, LD, and PO were also elevated over neuron levels. Acute drug exposure elucidated further cell-type specificity revealing a difference in the magnitude of responses and opposing trends in organelle signature metrics between cell types. Neurons responded more robustly with numerous significant differences in organelle signature metrics than astrocytes. Notably, many changes induced by drug exposures involved organelles and interactions that were significantly different between cell types in the control conditions. For example, neuron oxidative stress signatures exhibited increased MT-PO and ER-GL interactions, while astrocyte oxidative stress signatures showed reduced LS and increased LD amounts and interactions with other organelles.

The importance of mitochondrial function to neuron health has been robustly established^66,67^. MT are players in axon branching and neural circuitry^68–71^, calcium homeostasis for synaptic transmission^72–74^, and ATP production through OXPHOS and glycolysis^75,76^. Mechanistic studies have shown that certain mitochondrial functions, including OXPHOS, are essential to neuron survival^77,78^, while astrocytes can withstand inhibition of OXPHOS through upregulation of compensatory mechanisms^79,80^. MT interactions with ER and PO further this narrative as they are important to MT function. PO are key in intracellular ROS metabolism, and coupled interactions with MT may occur more frequently in neurons to modulate ROS levels from mitochondrial OXPHOS^81,82^. Interestingly, we noted increased MT-PO interactions during acute oxidative stress in neurons but not astrocytes. ER-MT membrane contact sites regulate mitochondrial fission, fusion, and mitophagy^6,83^ and maintain calcium homeostasis^84–87^. ER-MT interactions were the most common of all 15 interaction types across cell types, but the higher amount in neurons supports the idea that MT homeostasis is important to neuronal physiology.

LS play many important roles in astrocyte functions, including involvement in gliotransmission at neuronal synapses^88^, exocytosis of surface receptors during astrogliosis^89^, homeostatic phagocytosis of synapses^90^, and degradation of aberrant protein buildup in disease^91–94^. Qualitative observations of astrocytic LS in our dataset show the presence of larger LS than those observed in neurons. Secretory lysosomes, large LAMP1+ vesicular structures^95^, have been identified as the main exocytic vesicles involved in gliotransmission and astrogliosis^96,97^. Additionally, LS subcellular localization and interaction with other organelles suggest that astrocytes could have a different metabolic state than neurons in our culture system. Perinuclear clustering of LS has been linked to increased LS acidification^98^, LS fusion with autophagosomes for autophagy^99^, high cholesterol availability^100^, and additional phenotypes^101,102^. The presence of more numerous LS-GL interactions in astrocytes supports this idea, as LS-GL membrane contact sites mediated by tethering complexes, such as the RAB34-RILP-FLCN complex^45^, are related to the suppression of MTORC1 activity and the upregulation of autophagy^103,104^. We also noted higher LS-LD and LS-PO interactions. Membrane contacts between these organelles are involved in lipid metabolism^105–109^. Moreover, astrocytes showed a trending increase in LD volume upon acute stress. Others have found that astrocytes contain more LD in response to stress, possibly to protect against damaging lipid peroxidation^110–112^. Together, these findings suggest astrocytes in our culture system carry out more metabolic processes related to lipid metabolism and autophagy than neurons.

The importance of our findings to neuron and astrocyte physiology is further supported by genetic studies of neurodegenerative diseases. Mutations in MT-associated proteins result in neuronal MT dysfunction in AD, PD, ALS, and Huntington’s disease (HD)^113^. For instance, mutations in superoxide dismutase 1 (SOD1) cause 20% of familial ALS cases and lead to preferential accumulation of mutant SOD1 protein at MT in spinal cord motor neurons^114,115^. Huntingtin (HTT) repeat expansions, the mutant form of HTT (mHTT) responsible for HD, result in the mitochondrial permeability transition pore opening through the direct action of mHTT protein on the outer mitochondrial membrane^116^. mHTT deficits primarily occur by neuron-dependent mechanisms with lesser astrocyte-derived effects^117^. Furthermore, many lysosome storage disorders (LSDs) result in progressive neurodegeneration and implicate astrocytes as a main player in neuronal defects^118^. In one instance, astrocyte-specific deletion of the *SUMF1* gene implicated in multiple sulfatase deficiency (MSD) induced lysosomal storage and autophagy defects in astrocytes and concomitant degeneration of cortical neurons *in vivo*^119^.

Despite the high cell-type specificity of neuron and astrocyte organelle signatures, some metrics were stereotyped across cell types. The distribution of organelles displayed similar trends in perinuclear to peripheral localization: GL was perinuclear; ER, LS, and MT were central; and LD and PO were peripheral. However, these distribution phenotypes are likely relative to the 2D culture model and may be altered from *in vivo* conditions^120^. We also noted some similarities among the acute stress conditions. The largest magnitude change from baseline in all cells was related to GL fragmentation. An orthogonal machine learning (ML)-based approach in fixed induce pluripotent stem cell (iPSC) derived neurons also identified alterations in GL following a 30-minute, 400 μM sodium arsenite exposure^121^. Our data corroborate and build upon their findings by characterizing the change in GL pinpointed by their ML screening approach; our targeted metrics revealed that the previously observed altered GL morphology likely represents GL fragmentation. This phenotype is common and often reversible in many physiological and pathological conditions^122^, including neuron hyperexcitation^123^, DNA damage^124^, oxidative or ER stress^11^, and cancer^125^; it is also thought to be one of the earliest phenotypes in AD, suggesting oxidative and ER stress may be involved in early disease pathogenesis^126^. Lastly, we noted that most of the organelle signature changes following acute stress were organelle interaction metrics. This likely reflects the modulation of membrane contact sites through post-translational modifications of tether proteins^127,128^. In contrast, organelle biogenesis and turnover occur in longer time scales that require transcriptional regulation^129–131^. Extended sodium arsenite and thapsigargin exposures will, therefore, likely reveal novel organelle signature phenotypes reflective of transcriptional regulation.

The organelle signatures presented here characterize commonly utilized primary cell monocultures and, therefore, can act as an organelle reference for many studies, similar to the use of -omics datasets. However, *in vivo*, neurons and astrocytes directly and indirectly interact within a shared environment^132–135^. Feedback between cell types could impact their organelle signatures. This is exemplified in co-culture studies. One such study determined neuron hyperexcitation resulted in the release of neuronal peroxidated lipids that were then taken up by astrocytes^136^. Astrocytes resultantly upregulated beta-oxidation to break down the toxic lipid species. These findings suggest that signatures of lipid metabolism in astrocytes may be altered in a neuronal activity-dependent manner. Elucidating organelle signatures in co-cultures and *in vivo* models will be an exciting future direction for organelle signature analysis.

The 234 metrics included above only represent a fraction of the total possible measurements. We demonstrated the use of additional metrics included from the larger 1418 metric dataset in Figure 3 and hope that the full quantitative dataset will provide others with a resource to supplement their mechanistic or comparative studies in these cell types. The dataset also provides a foundation for hypothesis generation related to neuron and astrocyte physiology, cell specificity, and acute sodium arsenite and thapsigargin exposure. Moreover, we encourage members of the community to explore the imaging dataset further. We supplied raw, linear unmixed, and deconvolved Z-stack and timelapse images as well as segmentation results for Z-stack images in our BioImage Archive data repository (http://www.ebi.ac.uk/bioimage-archive; accession number S-BIAD1445). Additional quantitative analyses of the segmented organelles could be used to answer specific research questions not yet answered with our quantitative analysis. One such possibility would be the exploration of subclasses of organelles based on size, shape, interaction types, or distribution metrics presented here to tease out more cellular pathways and functions for future investigation. Machine learning techniques could also be a promising approach to uncover complex or multi-organelle phenotypes that are challenging to discern through standard mathematical approaches. Likewise, the raw and linear unmixed images allow for the use of novel unmixing and segmentation approaches or the quantitative exploration of intensity images, especially in the case of the timelapse data.

Beyond this study, organelle signature analysis can be applied broadly across cell types and drug or stress conditions. Comparisons between sets of cells involved in the same tissues or diseases are a promising avenue to advance our understanding of cell specificity. The drug response variability of neurons and astrocytes shown here exemplifies the need for this type of investigation.

Additionally, using the organelle signature to characterize disease-associated mutations is a powerful approach to exploring complex pathomechanisms and novel targets for therapeutic intervention. Disease-associated signatures then present an opportunity to test therapeutic responsiveness by monitoring system-wide markers of phenotypic improvement.

## Methods

### Primary neuron cultures

Primary cortical neurons were prepared from embryonic day 18 (E18) Sprague Dawley rats (Charles River Laboratories, strain 001) as previously described^35^. Briefly, dissociated cortical neurons were plated onto tissue culture dishes coated with 1mg/mL poly-L-lysine (PLL; Sigma Aldrich). Neurons were maintained at 37°C/5% CO_2_ and fed twice weekly in glial conditioned neurobasal media (Gibco 21130-049) supplemented with 2% B27 (Gibco 17504-044), 1% horse serum (Gibco 26050088), 2 mM Glutamax (Gibco 35050061), and 100 U/ml penicillin/streptomycin (Corning 30- 002-Cl). On DIV3, 1 μM FDU (Sigma F0503) was added during feeding. For live-cell imaging, neurons were plated at 75,000 cells/well into 4-well chambered glass slides (Cellvis C4-1.5H-N). For fixed cell microscopy and Western blotting experiments, neurons were plated at 150,000 cells/well into 12-well tissue culture plates (Corning 3513) with or without glass coverslips (Fisher Scientific 12- 545-81), respectively. For viability assays, 15,000 cells were plated per well of a 96-well plate (Thomas Scientific 1167V76).

### Primary astrocyte cultures

Cortical astrocytes were cultured as previously described with a slightly modified protocol^137,138^. Primary astrocytes were isolated from post-natal day 1-2 (P1-P2) Sprague Dawley rats (Charles River Laboratories, strain 001). Cortices from both sexes were microdissected and enzymatically digested with papain (10 U/brain; Worthington Biochemical, LK003176l) for 25 min at 37°C. The tissue was washed three times and gently aspirated to remove residual papain with astrocyte growth media (AGM), composed of DMEM high glucose (Corning, 15-013-CV0), heat-inactivated 10% FBS (VWR, 97068-085), 1X GlutaMAX (Gibco, 35050061), 1X penicillin/streptomycin (Corning, 30-002-CI), 5 µg/ml bovine insulin (Sigma-Aldrich, I6634), and 5 µg/ml N-acetyl-L-cysteine (Cayman Chemical, 20261). The tissue was triturated with three fire-polished Pasteur pipettes with progressively narrower bores in AGM and filtered through a 70-μm cell strainer (Falcon 352350).

Equal volumes of the cell suspension were plated onto 75-cm^2^ flasks (Flacon 354638) coated with 10 µg/ml of poly-D-lysine (PDL; Sigma A-003-E). Cells equivalent to the amount in one set of cortices were plated per flask. Flasks were maintained at 37°C/5% CO_2_, with a complete media exchange on DIV1 to remove debris. When astrocytes formed a confluent monolayer by DIV3, loosely adherent cells were physically dissociated by striking the flasks by hand three times in pre-warmed PBS. Cells were maintained for an additional 72 hr in AGM. To encourage more stellate morphology, astrocytes were passages into a serum-free growth medium. Cells were washed with PBS, lifted with 0.25% trypsin/EDTA (Corning, 25-053-CI), and pelleted at 125 RCF for 5 min. Cells were then washed with serum-free growth medium (NB+H), composed of phenol-free Neurobasal (Gibco, 12348017), 5 ng/mL HB-EGF (R&D Systems, 259-HE-050), 1X B27 supplement (Gibco, 17504044), 1X GlutaMAX (Gibco, 35050061), 1X penicillin/streptomycin (Corning, 30-002-CI), and then centrifuged. Counted cells were plated onto dishes freshly coated with 10 ug/mL fibronectin in PBS for 15 minutes at 37°C/5% CO_2_ and maintained in NB+H media until transfection. For live cell imaging experiments, astrocytes were plated in 8-well chamber glass slides (CellVis C8-1.5H-N) at 50,000 cells/well. For fixed cell microscopy and Western blotting experiments, astrocytes were plated at 100,000 cells/well into 24-well tissue culture plates (Corning 3526) with or without glass coverslips (Fisher Scientific 12-545-81), respectively.

### Multispectral Imaging of Organelles

#### Simultaneous Organelle Labeling

Primary neuron (day in vitro; DIV 15) or astrocyte (DIV 9) cultures were simultaneously labeled with genetically encoded and dye-based fluorescent markers for the following six organelles:

**Table 1:** Organelle Markers

| **Table 1: Organelle Markers** |  |  |  |  |  |
| --- | --- | --- | --- | --- | --- |
| **Organelle** | **Marker** | **Source** | **Method** | **Concentrations for Neurons** | **Concentrations for Astrocytes** |
| Lipid droplets | BODIPY665 | Thermo Fisher Scientific, B3932 | Dye | 0.4 µg/mL in media | 0.5 µg/mL in media |
| ER | mApple-Sec61 | Kind gift from Christopher Obara<sup>8</sup> | Plasmid Transfection | 0.7 µg | 0.15 µg |
| Peroxisomes | mOrange2-SKL | https://www.addgene.org/54596/ | Plasmid Transfection | 0.07 µg | 0.015 µg |
| Golgi | Venus-SiT | https://www.addgene.org/56336/ | Plasmid Transfection | 0.2 µg | 0.05 µg |
| Mitochondria | Mito-GFP | Kind gift from Angelika Rambold<sup>7</sup> | Plasmid Transfection | 0.15 µg | 0.015 µg |
| Lysosomes | Lamp1-mTurquoise | https://www.addgene.org/98828/ | Plasmid Transfection | 0.3 µg | 0.1 µg |

The labeling procedure was carried out 24 hours before the start of the imaging period. For multispectral imaging of cells with all six fluorescent labels (“6-label”), the five plasmids were transfected simultaneously using Lipofectamine 2000 (Invitrogen 52758) following a modified version of the manufacturer’s protocol. For one well of a 4-well chamber slide containing neurons, the indicated amount of each plasmid was added to one tube containing 50 uL of neurobasal media (Gibco 21130-049) without supplements (NB-/-). A 1:1 ratio of Lipofectamine 2000 to plasmid (μL:μg) was added to a second tube containing 50 μL of NB-/-. Tubes were vortexed for ∼5 seconds, and then the plasmid dilution was pipetted into the Lipofectamine dilution. The combined solution was vortexed again and incubated at room temperature for 15 minutes. A 100 μL aliquot of the DNA:Lipofectamine solution was added to 500 μL of neuronal media (neurobasal + supplements as specified above) in the well of cultured neurons. The dish was rocked gently to mix. Cells were incubated in the transfection solution for 40 minutes at 37°C/5% CO_2_. All media was removed and replaced with 500 μL of neuronal media with BODIPY665 at the above-indicated concentration. Cells were incubated at 37°C/5% CO_2_ until imaging. Astrocyte labeling in 8-well chamber slides followed the same protocol using the abovementioned astrocyte concentration.

Volumes were adjusted based on the surface area of the plate used. For single-label controls, each of the plasmids or dye was added individually; the same DNA:Lipofectamine ratio and dye concentrations were used.

#### Multispectral Imaging

Multispectral imaging was carried out as previously described^34^. Briefly, images were acquired on a Zeiss LSM880 laser scanning confocal microscope with a 34-channel GaAsP spectral detector and live cell incubation chamber (Carl Zeiss Microscopy). The 458 nm, 514 nm, 561 nm, and 633 nm lasers were used simultaneously to excite cells labeled with one or all of the six organelle markers. Emitted light was collected by the spectral detector in lambda mode, a linear array of 34 photomultiplier tubes (PMT), in 9.7 nm bins from 468 to 690 nm wavelengths, resulting in 26- intensity channels in the output image. A 63x/1.4 NA objective lens was used to image Z-stack and time-lapse images with an XYZ voxel scaling of 0.08 μm x 0.08 μm x 0.41 μm. Z-stacks were acquired at a single time point with a pixel density of 1688x1688 per slice. Slice number was determined on a cell-to-cell basis with the goal of imaging the entire cell from top to bottom. The time-lapse images included a single Z-plane (pixel density of 1124x1124) imaged with Definite Focus (Carl Zeiss Microscopy) every 5 seconds for 5 minutes (60 frames). All cells (DIV16 neurons or DIV10 astrocytes) were imaged live and maintained at 37°C/5% CO_2_ for up to 2 hours per dish while imaging.

#### Image Processing

Spectral images were first processed into 6-channel images using the linear unmixing algorithm from ZEN Black software version 2.3 (Carl Zeiss Microscopy). Briefly, control images that were labeled with one of the six organelle markers were imaged as described above. A region of fluorescence intensity from each of the images was used as a reference spectrum for that fluorophore. All six reference spectra were utilized as inputs in the linear unmixing algorithm. The same reference spectra were used for all images within an experimental replicate. New spectra were created for different labeling conditions and cell types. The linear unmixing process resulted in a six-channel image where each channel represents the fluorescence intensity derived from each of the six fluorophores.

Linear unmixed images were then deconvolved using Huygens Essential software version 23.04 (Scientific Volume Imaging, The Netherlands, http://svi.nl). Each of the six intensity channels was deconvolved independently using the same settings across all conditions with the Workflow Processor module. Excitation and emission maxima (Ex/Em max) were estimated for each fluorophore based on the lasers utilized during imaging and the reference spectra created for linear unmixing. The following settings were used:

| Channel | Ex/Em Max | Algorithm | Acuity | SNR | Iterations | Background |
| --- | --- | --- | --- | --- | --- | --- |
| Lipid droplet | 633/690 nm | Classic MLE | 41.1 | 5.6 | 38 | 399 |
| ER | 561/593 nm | Classic MLE | -19.76 | 11 | 26 | 50 |
| Golgi | 561/575 nm | Classic MLE | 19.33 | 1.5 | 34 | 340 |
| Lysosomes | 514/530 nm | Classic MLE | -49.14 | 6.5 | 26 | 264 |
| Mitochondria | 458/504 nm | Classic MLE | -49.83 | 5.4 | 26 | 52 |
| Peroxisomes | 458/477 nm | Classic MLE | 81.66 | 5.5 | 53 | 25 |

Unless otherwise indicated, example images in figures include single Z-plane crops from time series or Z-stack images after deconvolution.

### Organelle Signature Analysis

#### Image Segmentation

Instance segmentations were created from independent intensity channels in the deconvolved Z-stack images. Methods for segmentation are included and described in detail in our Python-based segmentation and analysis package, infer-subc version 1.0 (https://github.com/ndcn/infer-subc). They combine previously published segmentation methods optimized for organelles from the Allen Cell Segmenter “Classic Image Segmentation Workflow”^139^ and methods from the scikit-image Python package^140^. Semantic segmentation was performed first, followed by instance segmentation based on connectivity. Specifically, organelle objects found in the semantic segmentation were considered separate objects if they did not neighbor another object in any direction (X, Y, Z, or diagonally). Because of this, organelles within one voxel’s distance are considered the same object.

The cell mask and nucleus were inferred from a combination of all six organelle labels using similar methods. The nucleus was considered the absence of organelle fluorescence intensity in the central area of the soma or cell body. Astrocyte images usually contain multiple cells per field of view; a single cell with optimal labeling intensities was selected from each image for quantitative analysis. Neuron images only contained the somatodendritic region of a single neuron each. The ER, cell mask, and nucleus mask underwent semantic segmentation to produce a single object per image. This decision was made *a priori* based on the knowledge that each cell will only contain one ER, cell area, and nucleus.

The same segmentation settings were applied to all images. Qualitative checks were performed to ensure the accuracy of each segmentation output. If errors were found, the segmentations were manually edited. In neuron images, 20.55% of the segmentations were manually edited; in astrocyte images, 35.86 % were manually edited. Edits to the cell mask and nucleus comprise 69.53% of the manually edited segmentations.

#### Organelle Signature Quantification (1418 metrics)

Quantitative analysis of neuron and astrocyte organelle signatures is included and described in detail in infer-subc (https://github.com/ndcn/infer-subc/tree/main/notebooks). After segmentation, organelle objects and interactions between organelles, defined as the region of overlap between two organelle objects, were quantified for features of morphology (e.g., amount, size, and shape) and distribution. The morphology of the cell mask and the morphology and distribution of the nucleus mask were also quantified. All per-object measurements were summarized per cell, resulting in 1418 organelle signature metrics.

The full list of organelle signature metrics, metric definitions, and per-object quantification is included in our BioImage Archive data repository.

#### Morphology Measurements

Measurements of object morphology were collected to quantify the amount, size, and shape of each organelle and interaction site. We used the scikit-image skimage.measure.regionprops_table function^140^ including the following measurements for each object: label, centroid, bounding box, area (i.e., volume), equivalent diameter, extent, Euler number, solidity, and major axis length. The minimum intensity, maximum intensity, and mean intensity were also included for each channel in the cell mask quantification to allow for assessment of the fluorescence intensity of the organelle markers. Definitions of each property are included in sci-kitimage documentation here: https://scikit-image.org/docs/stable/api/skimage.measure.html#skimage.measure.regionprops.

The analyses are further summarized and were run utilizing infer_subc.utils.stats.get_org_morphology_3D and infer_subc.utils.stats.get_region_morphology_3D functions. All measurements were collected in “real-world” units (e.g., μm, nm), not voxel units.

#### Distribution Measurements

Distribution measurements were taken to assess the spread of organelles and interaction sites from the nucleus to the cell membrane (XY) and from the bottom to the top of the cell (Z).

Quantification of the XY distribution is based on the MeasureObjectIntensityDistribution module included in CellProfiler^141^. Briefly, a sum projection of each cell mask was created. Then, concentric rings proportional to the nucleus and cell shape were drawn, beginning at the edge of the nucleus mask and ending at the edge of the cell mask for each cell (Figure S4A). Each ring was utilized as a separate subcellular region from which we measured the volume of each organelle and interaction site from all Z planes. The centralmost ring included the nucleus and volumes above and below it, and the most peripheral ring included all the neurites in neurons or projections in astrocytes. To quantify the Z distribution, the bottom and top of the cell were defined by the lowest and highest Z planes that contained a portion of the cell mask, respectively. The Z planes were separated into ten approximately equal regions from which organelle volume was measured.

Each XY and Z region contained a different cell mask volume depending on the cell’s shape (Figure S4B-C). To control for this and differences in organelle amount per cell, the organelle, interaction site, and nucleus volume values were normalized to total object volume and region size. We represented the normalized volumes as histograms (Figure 4A-B, S4D). Histogram summary statistics were then used to summarize the spread of the organelles across the set of XY and Z regions more succinctly. To assess the variance in volume within each XY region, we also calculated the coefficient of variance per organelle and interaction site volume within each XY region.

Beginning at the center of the nucleus, radial lines were created to divide each XY region into eight equal fractions (Figure S4A, bottom panel red dashed lines). CV of the normalized volumes between each of the eight sections was calculated for each organelle and interaction site, and the median value per cell was included in the statistical analysis.

These analyses were run using infer_subc.utils.stats.get_XY_distribution and infer_subc.utils.stats.get_Z_distribution functions.

#### Batch Processing Quantification

Quantification for all images in each experiment was batch-processed using the infer_subc.utils.make_all_metrics_tables function. All experimental replicates were combined, and the data was summarized per cell using the infer_subc.utils.batch_summary_stats function. This produced the 1418 organelle signature metrics. Quantification for each experiment and the summary statistics per cell for the entire dataset are available in our repository on BioImage Archive.

#### Variable Curation (234 metrics)

To reduce redundancy and improve statistical power for the statistical comparisons included here, the 1418 organelle signature metrics were narrowed down to 234 metrics that were used to screen major differences between conditions in this study. The variables were chosen to increase the potential of seeing differences in organelles irrespective of the biological comparison being made. The values summarized per cell are described here:

| **Curated Organelle Signature Metrics (234)** |  |  |  |
| --- | --- | --- | --- |
| Organelles: ER, lysosomes (LS), mitochondria (MT), peroxisomes (PO), Golgi (GL), and lipid droplets (LD) |  |  |  |
| Interactions: the 15-pairwise interaction sites possible between each of the six organelles |  |  |  |
| Objects | Category | Name | Description |
| Organelles; interactions | Morphology | Count | The number of unique objects per cell. |
| Organelles; interactions | Morphology | Total volume | The sum of all unique object volumes (`skimage.measure.regionprops`) per cell. |
| Organelles; nucleus | Morphology | Volume Fraction | The proportion of the cell volume occupied by the object; calculated by dividing the total object volume by the cell mask volume (`skimage.measure.regionprops`). |
| Organelles; interactions | Morphology | Median Object Volume (not ER) | The median volume (`skimage.measure.regionprops`) of all unique objects per cell. |
| Organelles; interactions | Morphology | Standard Deviation of Object Volume (not ER) | The standard deviation of volume (`skimage.measure.regionprops`) of all unique objects per cell. |
| Organelles | Morphology | Median Object Surface Area to Volume Ratio | The median surface area to volume (`skimage.measure.regionprops`) ratio of all unique objects per cell.<br>Surface area was estimated from a 2D surface mesh of the 3D object (`skimage.measure.marching_cubes`) using `skimage.measure.mesh_surface_area`. The ratio was computed in `infer-subc`. |
| Organelles | Morphology | Median Object Equivalent Diameter | The median equivalent diameter (skimage.measure.regionprops) of all unique objects per cell.<br>Skimage defines the equivalent diameter as: “The diameter of a circle with the same area as the region.” In 3D, this equates to the diameter of a sphere with the same volume as the region. |
| Organelles | Morphology | Median Object Euler Number | The median Euler number (skimage.measure.regionprops) of all unique objects per cell.<br>Skimage defines Euler number as: “For 3D objects, the Euler number is obtained as the number of objects plus the number of holes, minus the number of tunnels, or loops.” |
| Organelles | Morphology | Median Solidity | The median solidity (skimage.measure.regionprops) of all unique objects per cell.<br>Skimage defines solidity as: “Ratio of pixels in the region to pixels of the convex hull image.” |
| Organelles | Morphology | Median Major Axis Length | The median major axis length (skimage.measure.regionprops) of all unique objects per cell.<br>Skimage defines major axis length as: “The length of the major axis of the ellipse that has the same normalized second central moments as the region.” |
| Organelles;<br>interactions;<br>nucleus | Distribution | XY Mode | The concentric XY region with the most normalized organelle volume per cell. Normalized volume values were utilized as the frequency distribution to generate a histogram, and the mode was calculated from the resulting histogram. |
| Organelles;<br>interactions;<br>nucleus | Distribution | XY Standard Deviation | The standard deviation (SD) of the normalized volumes across the five concentric XY regions. Normalized volume values were utilized as the frequency distribution to generate a histogram, and SD was calculated from the resulting histogram. |
| Organelles; interactions; nucleus | Distribution | Z Mode | The Z region with the most normalized organelle volume per cell. Normalized volume values were utilized as the frequency distribution to generate a histogram, and the mode was calculated from the resulting histogram. |
| Organelles; interactions; nucleus | Distribution | Z Standard Deviation | The standard deviation (SD) of the normalized volumes across the ten Z regions. Normalized volume values were utilized as the frequency distribution to generate a histogram, and SD was calculated from the resulting histogram. |
| Organelles; interactions; nucleus | Distribution | XY Median Coefficient of Variance per Bin | The coefficient of variance (CV) of the volumes per XY region was calculated and summarized as the median CV value per cell. |

#### Outliers

Outlier cells were considered based on the values of all 234 organelle signature metrics, not each metric individually. No outliers were removed from the dataset prior to statistical analysis.

#### Principle Component Analysis

Principle Component Analysis (PCA) was carried out using the GraphPad Prism 10.2.3 Windows, GraphPad Software, Boston, Massachusetts USA, www.graphpad.com. The curated list of 234 organelle signature metrics was first preprocessed by removing variables with a standard deviation of 0 and then standardized so that each variable had a mean of zero and a standard deviation of 1. Principle components (PCs) were selected using parallel analysis (1000 Monte Carol simulations; selection based on eigenvalues greater than the 95% percentile in simulation; random seed auto-selected).

#### Pairwise Comparisons

Pairwise comparisons between cell types and drug exposure conditions were made using a Mann-Whitney U test in GraphPad Prism 10.2.3 Windows, GraphPad Software, Boston, Massachusetts USA, www.graphpad.com. Significance was determined using the two-state linear step-up procedure of Benjamini, Kreiger, and Yekutieli with a False Discovery Rate (FDR) of 10% (q=0.1). All 234 organelle signature metrics were included in the analysis. However, metrics were removed if either comparison group contained one or fewer data points (e.g., only one cell or no cells have LD, so there were insufficient volume data points). Significance is denoted as follows: q > 0.1 (no significance), q ≤ 0.1 (*), q ≤ 0.05 (**), q ≤ 0.01 (***), and q ≤0.001 (****). The q-value and mean rank difference from this analysis, along with the number of cells (N), mean, median, minimum, maximum, standard deviation (SD), and percent coefficient of variance (%CV) for each experimental condition are summarized for neuron versus astrocyte and control versus drug exposure conditions in Tables S1 and S3, respectively.

Metrics outside of the curated analysis were analyzed independently. Specifically, the percent PO in LS-PO interactions (Figure 3D) utilized a Mann-Whitney U test from GraphPad Prism 10.2.3 Windows, GraphPad Software, Boston, Massachusetts USA, www.graphpad.com. P-value significance is denoted as follows: p > 0.05 (ns), p ≤ 0.05 (*), p ≤, .01 (**), p ≤ 0.001 (***), and p ≤ 0.0001 (****).

#### Correlation

The correlation of organelle signature metrics was based on Spearman’s correlation (r) and calculated along with p-values for significance in GraphPad Prism 10.2.3 Windows, GraphPad Software, Boston, Massachusetts USA, www.graphpad.com. P-value notation is summarized above. Spearman’s r and p-values for all correlation analyses are included in Table S2.

#### Hierarchical clustering

Hierarchical clustering was performed in R using the Pretty Heatmaps package version 1.0.12^142^. The list of organelle signature metrics processed for PCA (i.e., metrics with SD of 0 removed and data normalized) was used in this analysis. Cells were clustered using the ward.D method and Euclidian distance measurement.

### Cell Viability

To assess the impact of experimental conditions lasting more than 1-2 hours on primary cell viability, the CellTiter-Blue Cell Viability Assay (Promega G8080) was utilized following the manufacturer’s recommendations. In brief, primary cells were seeded in 96-well plates at the same density as imaging experiments as described above and prepared as if they were to undergo imaging. Instead of multispectral imaging at the 24-hour post-labeling timepoint, CellTiter-Blue reagent was diluted 1:5 into pre-warmed culture media and incubated on the cells for 4 hours at 37°C. Fluorescence (560_Ex_/590_Em_) was measured using a FLUstar Omega Plate Reader (GMB LabTech). Individual data points were background subtracted using the average fluorescence value from the “dead” negative control (70% EtOH exposure for 30 minutes) wells and normalized using the average fluorescence from the positive control (cells only) wells. Statistical comparison of conditions to the positive control (unmanipulated cells) was done using a repeat measures one-way ANOVA with a Geisser-Greenhouse correction and Dunnett’s test for multiple comparisons in GraphPad Prism 10.2.3 Windows, GraphPad Software, Boston, Massachusetts USA, www.graphpad.com. Neuron and astrocyte data were compared separately and consisted of two biological replicates with up to three technical replicates each.

### Cytotoxicity

To assess the level of cytotoxicity from experimental conditions that lasted 2 hours or less, CytoTox 96 Non-Radioactive Cytotoxicity Assay (Promega G1780) was utilized. The manufacturer’s protocol was followed. Briefly, three replicates of 50 μL of media from cultured cells were aliquoted into wells of a 96-well plate for all conditions tested. Media from unmanipulated cells of the same age were included as a negative control within each experiment. An equal volume of CytoTox 96 reagent was added and incubated in the dark at room temperature for 30 minutes. Stop solution was added to each well and bubbles were removed before reading the absorbance at 490nm. Percent cytotoxicity was calculated by dividing the average of the experimental replicates by the average of the “dead” cell control and multiplying by 100. Statistical comparison of conditions to the vehicle control was done using a one-way ANOVA with Dunnett’s test for multiple comparisons with a single pooled variance in GraphPad Prism 10.2.3 Windows, GraphPad Software, Boston, Massachusetts USA, www.graphpad.com.

### Western Blotting

Cells were lysed in ice-cold RIPA buffer (50mM Tris pH 8.0, 150mM NaCl, 1%NP-40, 5mM EDTA, 0.5% sodium deoxycholate, 0.1%SDS) containing additional inhibitors (2.5 mM β- glycerolphosphate (Sigma G9422) 2.5 mM NaF (Sigma 57920), 10 mM nicotinamide (Sigma 72340), 1 mM sodium orthovanadate (Alfa Aesar AA8110414), 1 mM phenylmethylsulfonyl fluoride (PMSF; Fisher Scientific 52-332), 1 uM Trichostatin A (Sigma T8552), 1X Protease Inhibitor Cocktail (Promega G6521), 2 uM okadaic acid (LC Labs O-2220). Protein samples were denatured in a mix of 6X sample buffer (Fisher Scientific 50-591-186) with dithiothreitol (DTT) and boiled at 98°C for 10 minutes before running on 12% Criterion™ TGX™ Precast Protein Gels (Bio-Rad 5671044). Samples were transferred to 0.2 um nitrocellulose membrane (Bio-Rad 162-0112) by wet transfer. Total protein amounts were assessed using PonceauS (Research Products International Corp P56200) protein stain and destained following manufacturer instructions. Membranes were blocked in 2% nonfat milk (Lab Scientific bioKEMIX 978-907-4243) in 1X TBS (Sigma T5912) for 30 minutes at room temperature while rocking. Primary antibodies were diluted into 2% milk and incubated overnight at 4°C while rocking. Membranes were washed three times in 1X TBST (1X TBS with 0.1% Tween 20; Sigma P9416), then incubated for one hour at room temperature while rocking in secondary antibody diluted in 2% milk. The membranes were washed again in the same way and then switched into 1X TBS for a final wash. Chemiluminescent or near-infrared immunofluorescence imaging was carried out using an ImageQuant LAS4000 machine or a Li-COR Odyssey CLx Infrared Imaging System (Li-COR Biosciences; Lincoln, NE), respectively. Phosphorylated eIF2α was detected with1:1000 rabbit anti-phospho-eIF2 (Cell Signaling CS3398) primary and 1:2000 goat anti-rabbit-HRP (Invitrogen 32460) secondary antibodies; GAPDH was detected with 1:500 chicken anti-GAPDH (Sigma AB2303) primary and 1:10,000 goat anti-chicken IRDye 800CW (LI-COR 926- 32211) secondary antibodies.

### Stress Granule Imaging

Primary neurons or astrocytes cultured on glass coverslips at the same plating density as was used for live cell imaging experiments were fixed for 10 minutes at 37°C/5% CO_2_ in a solution containing 2% paraformaldehyde (PFA; Electron Microscopy Sciences 15713) and 0.05% glutaraldehyde in PHEM buffer (EMS, 11162) prewarmed to 37°C. Coverslips were rinsed three times in PBS (Corning 3822009) and permeabilized in 0.2% Triton X-100 (Sigma X100) for 8 minutes at room temperature. Coverslips were rinsed again in PBS, incubated in 2% milk for 1 hour at room temperature, and then incubated in primary antibody solution overnight at 4°C in a humid chamber. Two markers of stress granules, G3BP1 and TIA1, were used for detection: 1:3000 rabbit anti-G3BP1 (Protein Tech T3057- 2-AP, lot 000-5872) and 1:250 goat anti-TIA1 (Santa Cruz SC-1751, lot E1314). Coverslips were rinsed again in PBS, incubated in secondary antibody diluted in 2% milk for 1 hour at room temperature in the dark, rinsed again, and mounted to slides using SlowFade Mounting Medium with DAPI (ThermoFisher S36973) and sealant (Biotum 23005). The following secondary antibodies were used serially: staining round 1 used 1:500 donkey anti-goat-AF488 (LifeTech A11055, lot 1627966) followed by 1:500 goat anti-rabbit-AF568 (Invitrogen A11011, lot 2088069). After curing for at least 24 hours at room temperature, slides were imaged. Images were collected with a Zeiss LSM800 point scanning confocal microscope (Carl Zeiss Microscopy) using a 63X/1.4 NA oil immersion objective with 405 nm, 488 nm, and 561 nm lasers. The percentage of cells that contained stress granules was counted manually. Images were processed for visualization in ImageJ Fiji^143^. Example images are maximum intensity projections.

### Summary diagrams

The summary diagrams in Figures 1A, 6C, and S4A were created in Biorender: Figure 1A: Created in BioRender. Cohen, S. (2024) https://BioRender.com/t11q270 Figure 6C: Created in BioRender. Cohen, S. (2024) https://BioRender.com/b84o183 Figure S4A: Created in BioRender. Cohen, S. (2024) https://BioRender.com/z12e684

### Graphing and data visualization

Box plots, heatmaps, and dendrograms were created using R coding language^144^. Box plots and heatmaps were created with ggplot2^145^. Box plots display the median, range, and interquartile range. Individual data points represent per-cell values. Dendrograms were generated with Pretty Heatmaps as described above^142^.

PC scores scatter plots, volcano plots, bar charts, dot plots, and pie charts were created in GraphPad Prism 10.2.3 Windows, GraphPad Software, Boston, Massachusetts USA, www.graphpad.com.

### Data storage and availability

All imaging and quantitative data is available at BioImage Archive accession number S-BIAD1445^146^.

## Acknowledgments

Multispectral microscopy was performed at the UNC Hooker Imaging Core Facility, supported in part by P30 CA016086 Cancer Center Core Support Grant to the UNC Lineberger Comprehensive Cancer Center. We thank Wendy Salmon (Hooker Imaging Core) for expert imaging advice. This work was supported by the National Institutes of Health under awards T32 NS007431 (SR), F31 AG079622 (SR), R01NS105981 (TJC), and R35GM133460 (SC), and by Chan Zuckerberg Initiative award A23-0264-001 (SC).

## Supplemental File Legends

Table S1: **Neuron vs astrocyte organelle signature metric summaries and statistical comparison.** A table of the number of cells (N), mean, median, minimum, maximum, standard deviation (SD), and the q-value and mean rank difference from the Mann-Whitney U test for control neuron and astrocyte data. The 234 curated organelle signature metrics are included.

Table S2: **Correlation matrix of all organelle signature metrics and cell morphometrics for control neurons and astrocytes.** Spearman’s r and p-values for the correlation of all organelle signature metrics (234) and morphometrics for control neurons and astrocytes.

Table S3: **Treatment organelle signature metric summaries and statistical comparison.** A table of the number of cells (N), mean, median, minimum, maximum, standard deviation (SD), and the q- value and mean rank difference from the Mann-Whitney U test for sodium arsenite and thapsigargin treated neuron and astrocyte data. Comparison data summarizes the differences between treatment and control conditions for each cell type separately. The 234 curated organelle signature metrics are included.

**Movie S1: Multispectral imaging of organelles in live primary neurons & astrocytes**. Representative time-lapse images of control neurons and astrocytes labeled with fluorescent markers for the mitochondria (green), peroxisomes (orange), Golgi (yellow), lysosomes (cyan), lipid droplets (magenta), and ER (red). The aforementioned order of organelles corresponds to the order in which each channel appears in the movie. 2D snapshots were taken every 5 seconds for 5 minutes (60 frames).

**Movie S2: PO-MT interactions in control neurons and astrocytes.** Representative time-lapse images of control neurons and astrocytes captured by multispectral imaging. Peroxisome (magenta) and mitochondria (green) intensity channels are displayed. 2D snapshots were taken every 5 seconds for 5 minutes (60 frames).

**Movie S3: GL-LS interactions in control neurons and astrocytes.** Representative time-lapse images of control neurons and astrocytes captured by multispectral imaging. Golgi (magenta) and lysosomes (green) intensity channels are displayed. 2D snapshots were taken every 5 seconds for 5 minutes (60 frames).

**Figure S1.**
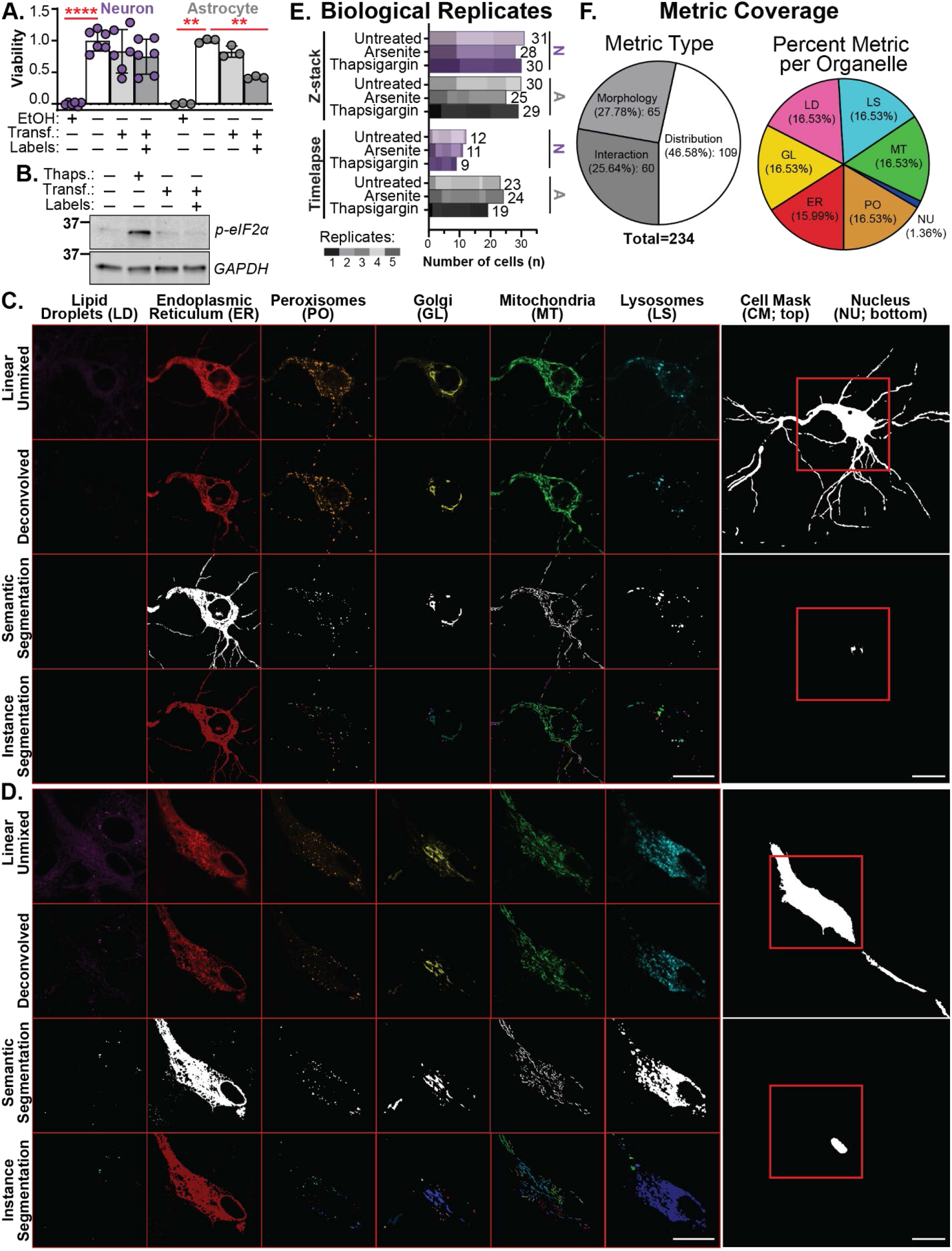
Multispectral imaging controls and segmentation outputs. A. Cell viability following multispectral imaging sample preparation (n=2). Bars represent the mean ± SD of the normalized CellTiter-Blue intensity measurement; individual data points represent experimental replicates. B. Representative western blot results of phosphorylated eIF2α levels following multispectral imaging preparation (n=3). C-D. Representative deconvolution and segmentation processing output for the example neuron (C) and astrocyte (D) included in Figure 1B-C; scale bars represent 20 μm. E. The number of cells analyzed across each biological replicate in the Z-stack organelle signature and timelapse datasets; neuron data (N) is shown in purple bars, and astrocyte data (A) is shown in grey bars. F. Percent coverage of metric types and organelles in the 234 curated organelle signature metrics.

**Figure S2.**
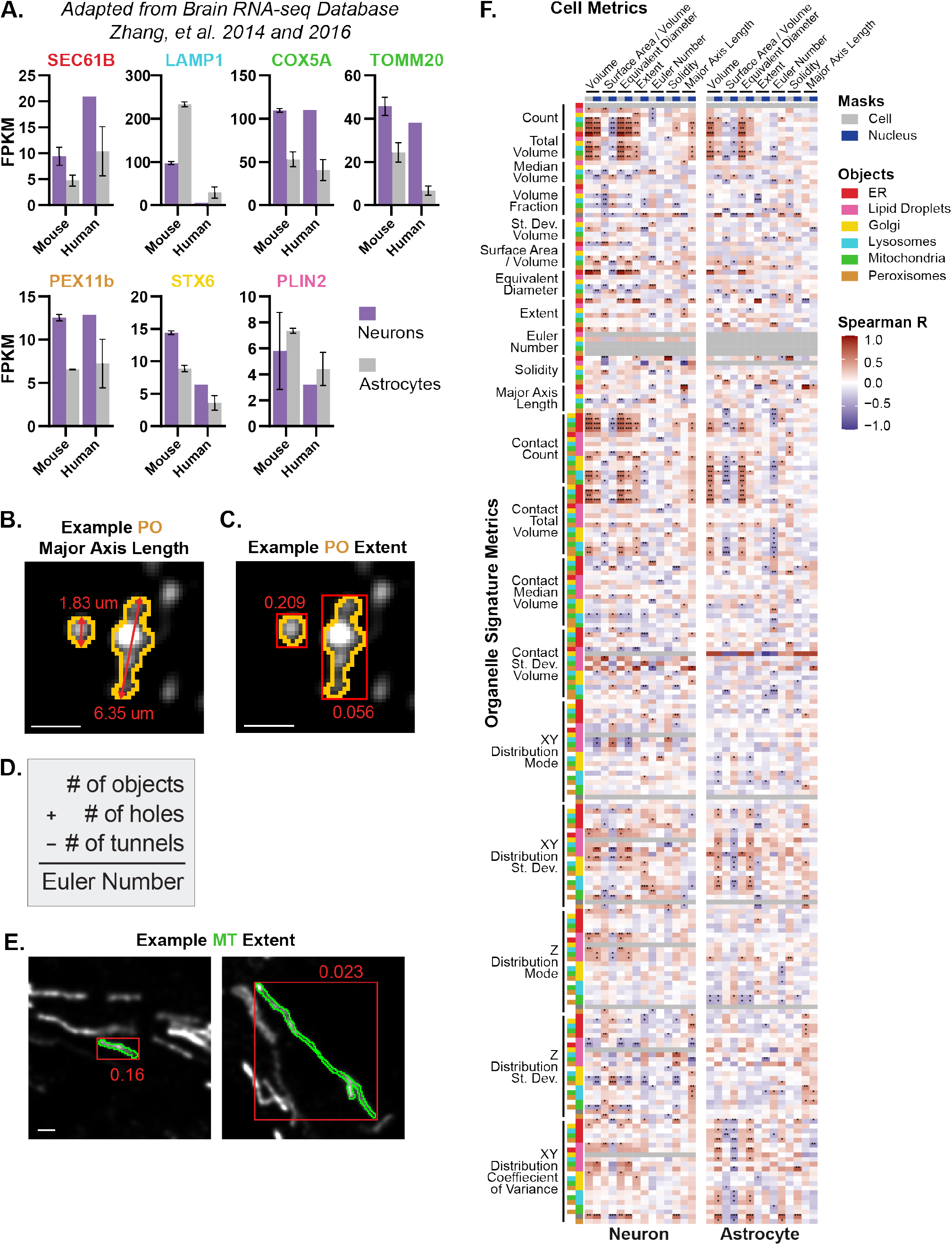
Neuron and astrocyte organelle morphology correlate with cell function and some cell shape metrics. A. RNA-seq results adapted from the Brain RNA-seq Database of stereotypical organelle markers; ER (SEC61B), LS (LAMP1), MT (COX5A and TOMM20), PO (PEX11b), GL (STX6), and LD (PLIN2). B, C, E. Example intensity images illustrate shape measurements of individual organelle objects (outlines); scale bars are 1 μm. D. Calculation for Euler number shape metric. F. Correlation heatmap of organelle signature metrics versus metrics of cell size and shape; asterisks represent p-values. Also, see Table S2.

**Figure S3.**
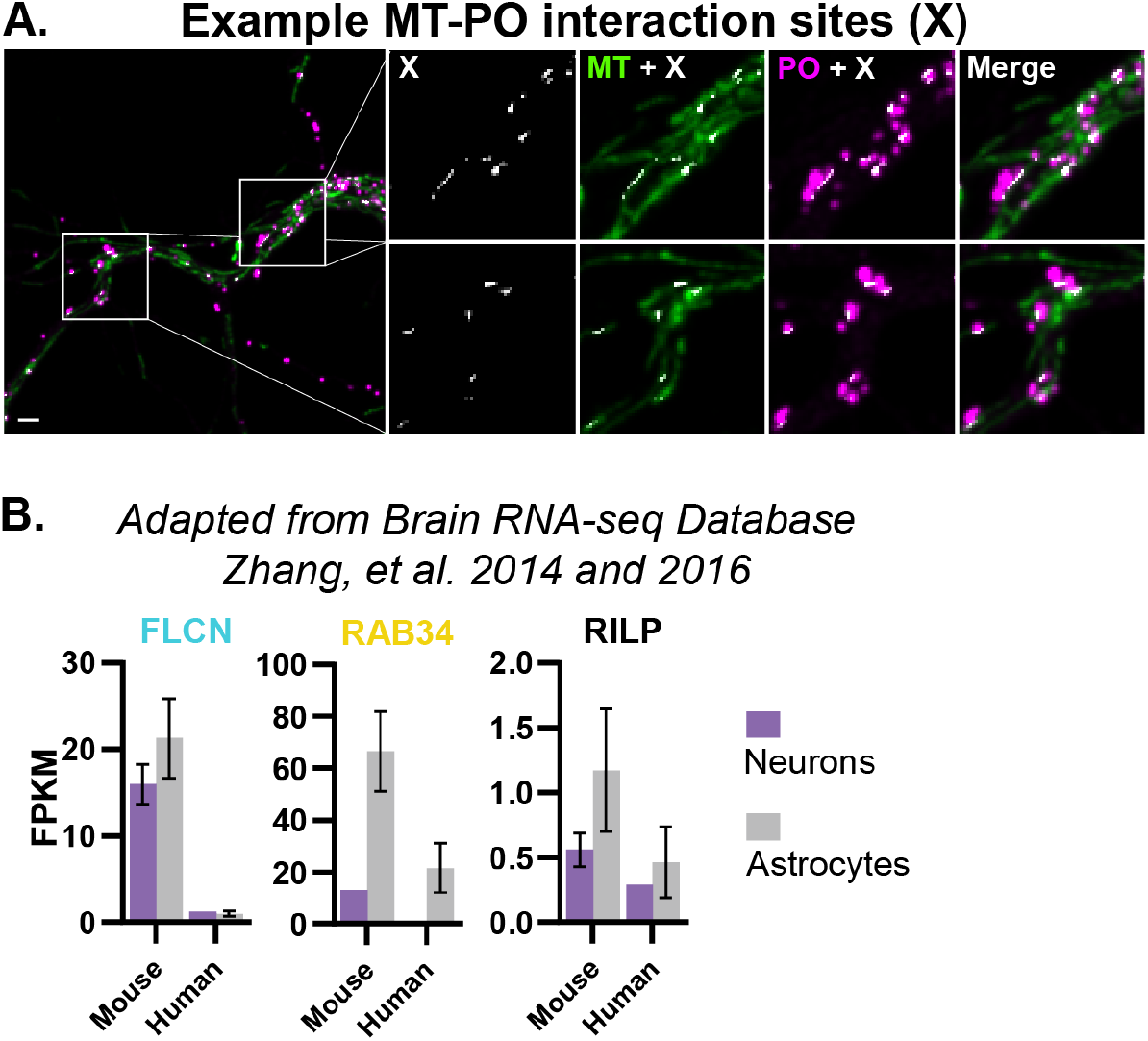
Organelle contact tethers trend with organelle interaction phenotypes. A. Example segmented MT-PO interaction sites (X; white) in a neuron overlayed onto the MT and PO intensity channels; scale bar is 1 μm. B. RNA-seq results adapted from the Brain RNA-seq Database of LS-GL membrane contact tethers; FLCN (LS localized), RAB34 (GL localized), RILP (tether between RAB34 and FLCN).

**Figure S4.**
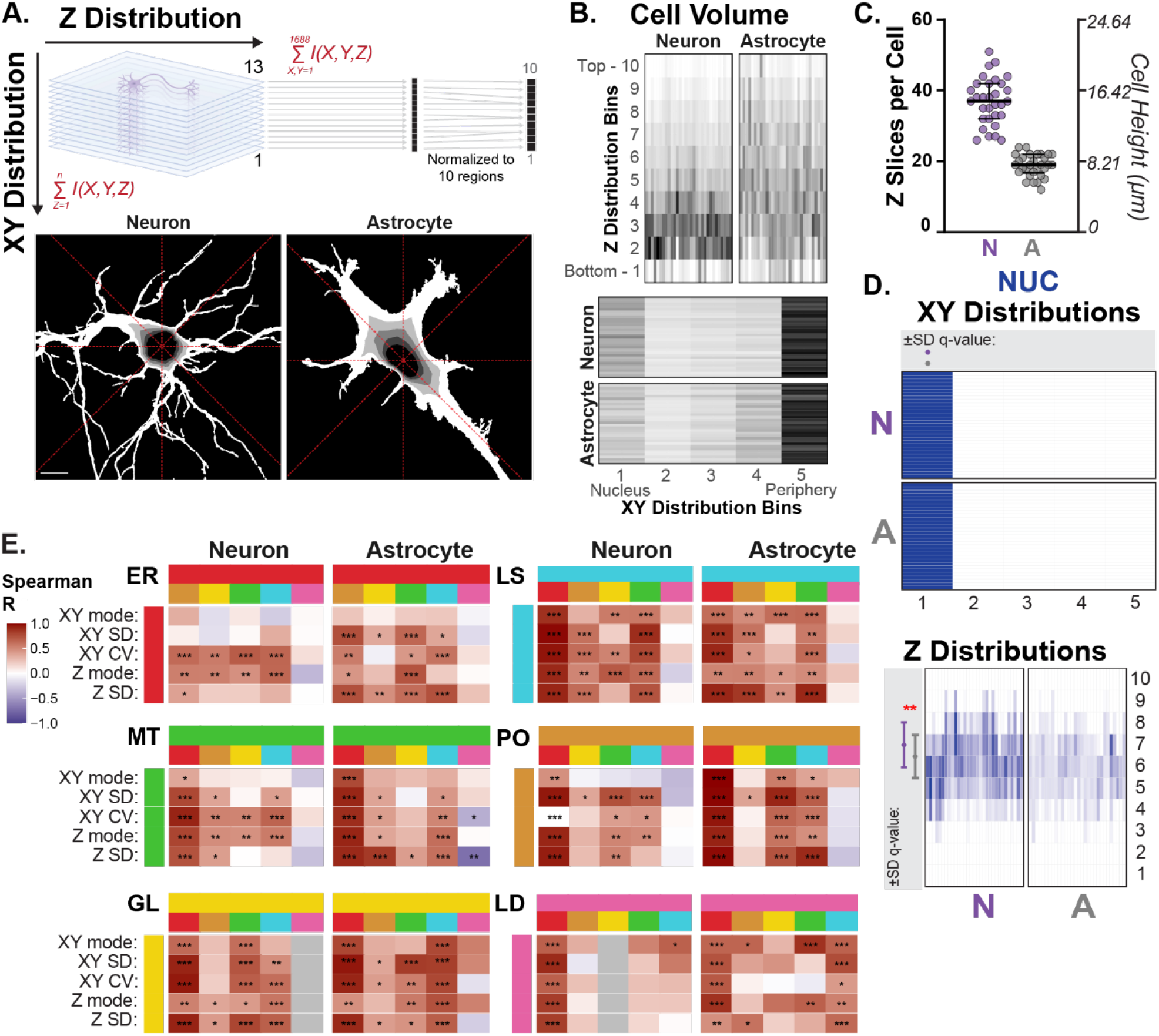
**Subcellular distribution of organelles and organelle interaction sites**. A. Schematic of XY and Z distribution measurements. To measure the XY distribution, segmented Z-stack images were made into sum projections where the number of voxels containing an object (intensity; I = 1) are summed together for each XY position across all Z-planes (Z=1 → n). Using the cell and nucleus mask sum projections, five XY regions were partitioned from the edge of the nucleus to the edge of the cell. The object volume per region is calculated. A, bottom: Greyscale images display example XY regions; red dashed lines denote the eight equal radial sections used to calculate coefficient of variance (CV) values for each XY region; scale bar is 10 μm. To measure the Z distribution, segmented Z-stack images were made into sum projections where the number of voxels containing an object (I = 1) are summed together for each Z-plane across all XY positions (X,Y=1 → 1688). A, top: Z-planes are divided into ten equal regions. B. Heatmaps of the total cell volume per XY (bottom) and Z (top) regions for all untreated control cells in the dataset; each column (Z) or row (XY) represents the cell volume per region for individual cells. C. Median and interquartile range of neuron and astrocyte cell height; data points represent individual cell values. D. Heatmaps of the normalized nuclei volumes per XY and Z regions. Grey boxes summarize the mode (data point; asterisk perpendicular to the bars) and SD (error bars; asterisk parallel to the bars) distribution metrics. E. Correlation heatmap of organelle interaction distribution metrics (x-axis) compared to their constituent organelle distribution metrics (y-axis); each row represents one type of distribution metric; grey boxes indicate correlation was not calculated; asterisks represent correlation p-values.

**Figure S5.**
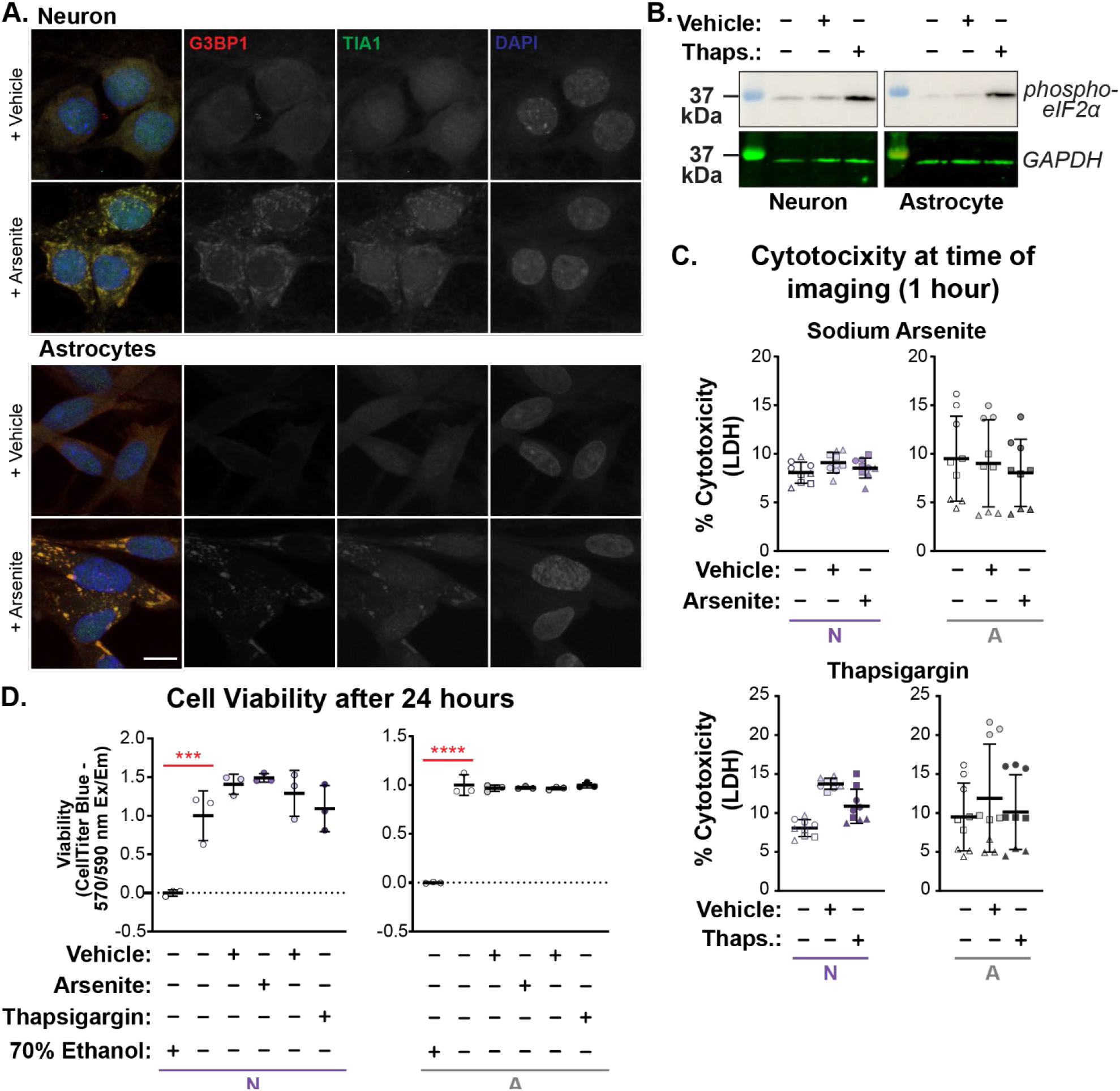
Sodium arsenite and thapsigargin exposure induce canonical oxidative and ER stress responses without a concomitant reduction in cell viability. A. Immunofluorescence microscopy of G3BP1/TIA1+ stress granules in neurons and astrocytes treated with 50 uM sodium arsenite for 1 hour; neurons: n=3 (43 cells), astrocytes: n=4 (37 cells); scale bar is 10 μm. B. Western blot of phospho-eIF2α in neurons and astrocytes treated with 25 nM thapsigargin for 1 hour; n=3. C. Mean ± SD of the percent cytotoxicity as measured by lactate dehydrogenase (LDH) levels following 1 hour of drug or vehicle exposure; asterisks denote significance as determined by one-way ANOVA; data points represent experimental replicates across biological replicates (shapes). D. Mean ± SD of the cell viability as measured by CellTiter Blue fluorescence 24 hours after exposing the cells to sodium arsenite or thapsigargin for 1 hour; asterisks denote significant differences from untreated controls as determined by one-way ANOVA; data points represent experimental replicates across biological replicates (shapes).

**Figure S6.**
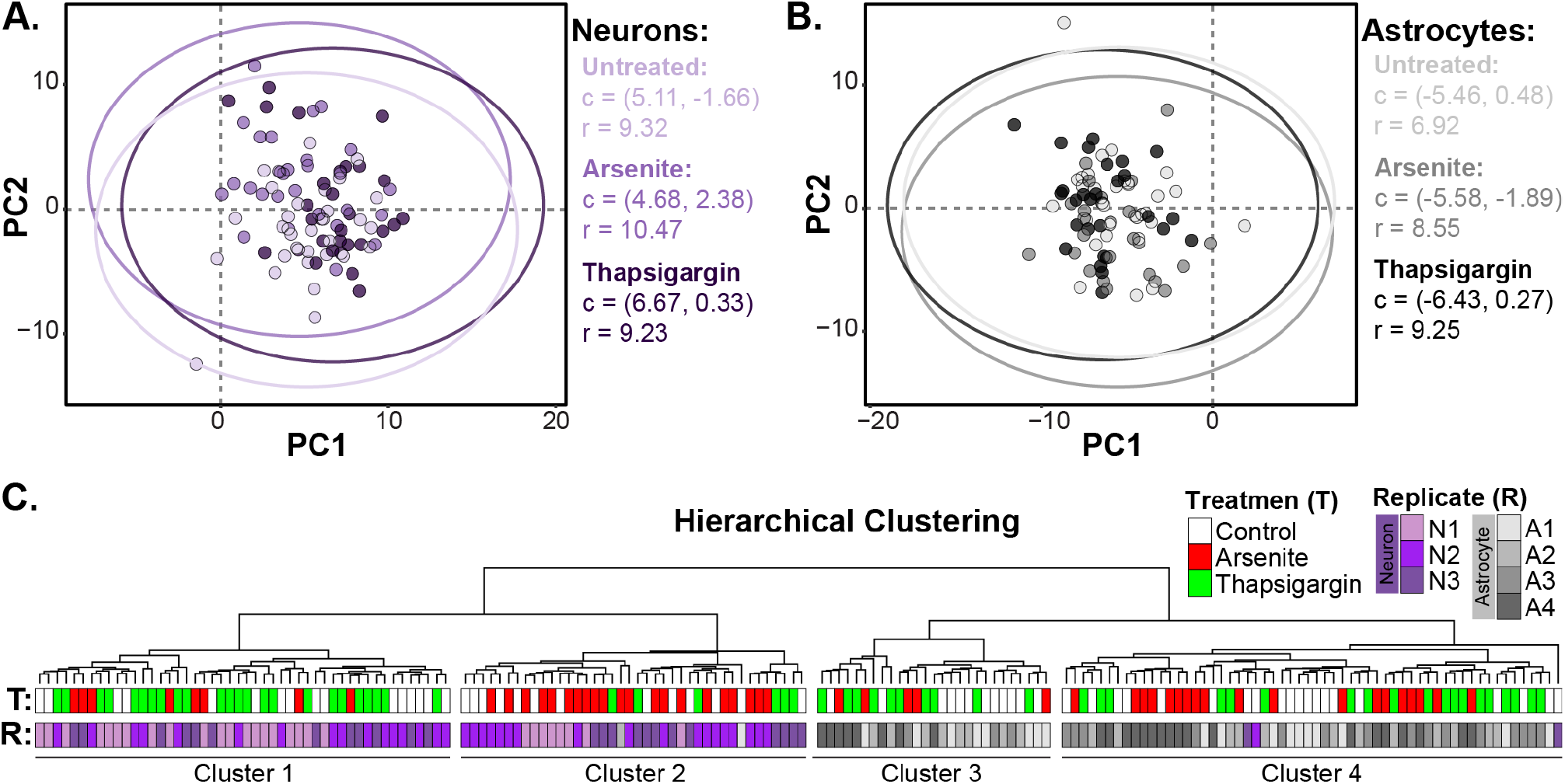
Stress-induced organelle modifications are subtle compared to cell type differences. A-B. PCA data from Figure 6A separated by cell type; circles summarize the center of the data points for each condition (c) and the spread of the data points as displayed by ±1.5 times the interquartile range (r). C. Hierarchical clustering of all neuron and astrocyte data across conditions revealed discernable clusters of neurons and astrocytes with three cells from neuron cultures and two cells from astrocyte cultures clustered with the opposite cell type; experimental replicates (R) are displayed as gradients of purple and grey; treatment conditions (T) are white, green, or red.

